# Genomic 8-oxoguanine is associated with transcriptionally active chromatin and elevated gene expression in *Plasmodium falciparum*

**DOI:** 10.64898/2026.06.21.733641

**Authors:** Dimple Acharya, Shruthi Sridhar Vembar

## Abstract

Epigenetic regulation is central to the developmental progression and pathogenicity of the unicellular eukaryotic parasite *Plasmodium falciparum*; yet, the contribution of DNA base modifications remains poorly understood. One such modification, 8-oxoguanine (8-oxoG), which was initially identified as an oxidative lesion and a marker of DNA damage, has since emerged as a transcriptional regulator in advanced eukaryotes. Given that *P. falciparum* encounters a highly oxidative environment in human blood, we investigated the potential gene regulatory role of 8-oxoG during its intra-erythrocytic developmental cycle (IDC). Using immunodetection assays, we first confirmed the presence of 8-oxoG in *P. falciparum* genomic DNA and observed a gradual increase in 8-oxoG abundance from ring to schizont stages. We then optimized oxidative DNA immunoprecipitation sequencing (OxiDIP-seq) for the highly AT-rich parasite genome and generated genome-wide 8-oxoG profiles across four IDC timepoints, which revealed reproducible enrichment of 8-oxoG at discrete genomic loci, with more than 50% of the peaks stable across developmental stages. Notably, 8-oxoG accumulated at putative G-quadruplex-forming sequences in the parasite genome and preferentially localized within exonic regions of protein-coding genes, exhibiting a marked enrichment near STOP codons and within 3′ untranslated regions. This in turn correlated with significantly higher steady-state transcript levels of 8-oxoG-marked genes, with stage-specific changes in 8-oxoG enrichment closely matching transcriptional activity. Furthermore, 8-oxoG-marked loci were preferentially associated with active and poised histone post-translational modifications, while showing no evidence of altered nucleosome occupancy. Collectively, these findings demonstrate that 8-oxoG is a widespread and non-random DNA modification in *P. falciparum* and suggest that it may function as an epigenetic mark associated with transcriptionally permissive chromatin and gene activation during parasite blood-stage development.

## Introduction

Decades of research have established that gene regulatory mechanisms, including epigenetic, transcriptional, post-transcriptional, translational, and post-translational processes, control the life cycle of the malaria parasite *Plasmodium falciparum* inside the human host and mosquito vector (Batugedara et al., 2017; Cui et al., 2015a; Doerig et al., 2015; Hughes et al., 2010; Voss et al., 2014; M. Zhang et al., 2013). By activating or silencing specific sets of genes in a tightly regulated fashion, these mechanisms are responsible for the distinct physiological and morphological characteristics of each *P. falciparum* developmental stage (Bozdech et al., 2003a; López-Barragán et al., 2011; van Biljon et al., 2019; Young et al., 2005). Specifically, histone post-translational modifications (PTMs), together with histone variants, non-coding RNAs (ncRNAs), chromatin remodellers and nuclear architecture, are now recognized as central regulators of parasite virulence, host adaptation, developmental transitions and transmission (Coetzee et al., 2017; Comeaux & Duraisingh, 2007; Connacher et al., 2021; Crowley et al., 2011; Gómez-Díaz et al., 2017; Gupta et al., 2017; Gupta & Bozdech, 2017; Herrera-Solorio et al., 2019; Issar et al., 2009; Kaur et al., 2016; M. Kumar et al., 2021; Ngwa et al., 2017, 2021; Saraf et al., 2016; Trelle et al., 2009; Witmer et al., 2020). Yet, the role of DNA base modifications in epigenetically regulating *P. falciparum* gene expression is relatively under-studied.

During the parasite’s intra-erythrocytic developmental cycle (IDC), attempts to detect and functionally characterize 5-methylcytosine (5mC), the best studied eukaryotic DNA base modification with an essential epigenetic role in advanced eukaryotes (Boland et al., 2014; Breiling & Lyko, 2015; Hack et al., 2016; Klose & Bird, 2006; S. Kumar et al., 2018; Sun et al., 2014), were inconclusive (Gissot et al., 2008; Hammam et al., 2020; Lucky et al., 2023; Ponts et al., 2013), possibly due to the low G+C content (∼19.4%) of the *P. falciparum* genome (Gardner et al., 2002a). For instance, we and others reported low levels of 5mC in the *P. falciparum* genome (Hammam et al., 2020; Lucky et al., 2023), contradicting a report from the Le Roch group that detected >1% of cytosines as being modified to 5mC during *P. falciparum* red blood cell (RBC) stages (Ponts et al., 2013); instead, our data supported the existence of high levels of a modification that had a similar mass to the oxidative derivative of 5mC, 5-hydroxymethylcytosine (5hmC) (Hammam et al., 2020). We further showed that the enrichment of this 5hmC-like modification within exons of protein-coding genes correlated with increased steady state transcript levels, indicating a possible role for this modification in epigenetic regulation (Hammam et al., 2020). Nonetheless, the exact nature of the 5hmC-like modification in *P. falciparum* is still unknown, and other DNA base modifications remain poorly characterized. One such modification is 8-oxoguanine (8-oxoG).

Among the four DNA bases, guanine is the most susceptible to oxidative damage (Dizdaroglu, 2002), frequently forming 8-oxoG due to its relatively low oxidation potential (Neeley & Essigmann, 2006). This lesion is moderately mutagenic and, if left unrepaired, can contribute to genomic instability, cellular transformation, and the initiation of cancer (Hegde et al., 2008). In mammalian cells, 8-oxoG is typically recognized and excised by the DNA glycosylase/AP lyase OGG1 through the base excision repair (BER) pathway (Boiteux & Radicella, 2000). Notably, beyond its role in DNA repair, emerging evidence suggests that 8-oxoG also participates in gene regulation and telomere end maintenance in response to reactive oxygen species (ROS) accumulation (Fedeles, 2017; Zhong et al., 2024). Specifically, its formation within G-quadruplex structures of gene regulatory regions has been associated with the epigenetic modulation of oxidative stress response genes (Ba & Boldogh, 2018a; Pan et al., 2016a), wherein, chromatin remodelling and transcriptional activation is facilitated by recruiting OGG1 to sites of oxidative damage. Other reports have further established the 8-oxoG and OGG1 axis as a regulator of transcriptional outcomes in mammalian cells (Allgayer et al., 2016; Ba & Boldogh, 2018a; Fleming, Ding, et al., 2017; Pan, Hao, et al., 2023a; Sampath & Lloyd, 2019). Mechanistically, the formation of 8-oxoG within promoter or regulatory regions serves as a signal for the recruitment of OGG1, which binds to the oxidized base with high affinity (Ba et al., 2014a). Under conditions where its glycosylase activity is transiently inhibited or delayed, OGG1 remains bound to DNA and functions as a structural and regulatory factor rather than simply initiating the BER pathway, facilitating local chromatin remodelling by inducing DNA bending and destabilization, thereby increasing accessibility of regulatory regions (Fleming, Ding, et al., 2017; Gillespie & Wilson, 2007; Pan et al., 2016a, 2017; Pastukh et al., 2015; Visnes et al., 2018).

Given that during human blood stage infection, *P. falciparum* develops within the highly oxidizing environment of the RBC (Becker et al., 2004), we set out to explore whether the parasite relies on 8-oxoG accumulation in gene regulatory regions to epigenetically activate transcription. This could in turn explain how the parasite integrates oxidative cues to fine-tune transcriptional responses, contributing to its survival and pathogenicity within human blood. Because the presence of 8-oxoG has not been reported in *P. falciparum* during its IDC, although a putative homolog of OGG1 encoded by the *PF3D7_0917100* gene has been partially characterized (Tiwari et al., 2024), we first confirmed the presence of 8-oxoG in *P. falciparum* genomic DNA and analysed its genome-wide distribution patterns at four distinct timepoints of the IDC using DNA immunoprecipitation with anti-8-oxoG antibodies followed by high throughput sequencing (OxiDIP-seq). This revealed that 8-oxoG-containing peaks are enriched within gene bodies, with a strong peak close to the STOP codon and within the 3’ untranslated region (UTR), and correlate with elevated steady state transcript levels during the IDC. Moreover, genomic 8-oxoG enrichment appears to be associated with a subset of activating/poised histone PTMs, as well as G-quadruplex-forming sequences, in keeping with studies in human cell lines. Overall, our data point toward a transcriptional activating role for 8-oxoG in *P. falciparum* RBC stages.

## Materials and Methods

### In vitro culturing of P. falciparum

Asexual blood stages of the *P. falciparum* laboratory strain 3D7 were grown according to Trager and Jensen (Trager & Jensen, 1976) with a few changes. Briefly, a mixed stage culture of *P. falciparum* was grown in O+ human blood at a hematocrit of 4% in Roswell Park Memorial Institute (RPMI) 1640 medium containing L-glutamine (Invitrogen) supplemented with 1% v/v Albumax II (Invitrogen) and 0.1 mM hypoxanthine at final concentration and 10 mg/ml gentamicin (ThermoFisher Scientific). The cultures were grown in a gas environment of 5% CO_2_ at 37°C, to a parasitemia of 3–8% before harvesting for downstream analysis. Parasite development was monitored by Giemsa staining of blood smears. Parasites were synchronized by 5% D-Sorbitol (Sigma) lysis at ring stages (Lambros C, Vanderberg JP, 1979).

### Genomic DNA isolation

Infected human erythrocyte cultures synchronized at the ring (8–12 h post-invasion (hpi)), trophozoite (28–32 hpi) or schizont (40–44 hpi) stages of the *P. falciparum* 3D7 strain were harvested and free parasites obtained by saponin lysis (Ljungström et al., 2008). Genomic DNA (gDNA) was prepared using the DNeasy Blood and Tissue kit (Qiagen) as per manufacturer’s instructions and treated with RNase to remove any contaminating RNA molecules. Additionally, gDNA were isolated from human white blood cells (hWBC), HEK293 cells, *Saccharomyces cerevisiae* (*S. cerevisiae*), and *Escherichia coli* (*E. coli*) using the GeneJET Genomic DNA Purification Kit (Thermo Fisher Scientific). DNA concentration and quality were measured using the NanoDrop^TM^ Microvolume Spectrophotometer (Thermo Fisher Scientific) and Qubit Fluorimeter (Thermo Fisher Scientific).

### Quantification of 8-oxodG using South-western blotting

South-western blotting assays of *P. falciparum* gDNA were performed as described (Bowen et al., 1980) but without using a DNA-binding protein. Purified gDNA prepared as described above was denatured at 95℃ for 10 min and chilled on ice for 5 min. Different amounts of gDNA (400, 200, 100, 50, and 25 ng), were spotted onto a piece of Hybond-N + membrane (RPN303B, GE healthcare, UK) and then UV cross-linked at 70000 μJ/cm^2^ for 1 min. The membranes were blocked with 5% milk in phosphate-buffered saline (PBS) + Tween-20 (PBST), and probed using mouse monoclonal antibodies against 8-oxoG (MAB3560, Millipore) at 1:1000 dilution overnight in the cold room. For signal amplification, the membrane was incubated with horse radish peroxidase (HRP)-conjugated anti-mouse secondary antibodies (CST-7076, Cell Signalling Technologies) at 1:5000 dilution in PBST for 1 h at room temperature. Signals were detected using ECL Western Blotting Detection Reagents Clarity kit (Bio-Rad) and captured using the ChemiDoc Imaging System (Bio-Rad).

### OxiDIP-seq

OxiDIP-seq, adapted from Fleming et al. (Fleming, Zhu, et al., 2017), was used to determine the 8-oxoG genome-wide distribution at different time points of the *P. falciparum* IDC. Briefly, synchronized parasites at the early ring (8-10 hpi), late ring (18-20 hpi), trophozoite (28-30 hpi) and schizont (38-40 hpi) stages of the *P. falciparum* 3D7 strain were harvested and gDNA isolated as described above for three biological replicates. 2 μg of gDNA per immunoprecipitation reaction was sonicated in 50 μl TE buffer (100 mM Tris–HCl pH 8.0, 0.5 M EDTA pH 8.0) to generate random fragments ranging in size between 200 and 600 bp using a Bath Ultrasonicator (30” ON and 90” OFF for 20 cycles). 2 μg of fragmented gDNA in 50 μl TE Buffer was incubated with 50 μl Dynabeads Protein G (Cat. No. L00274, Genscript; previously saturated with 0.5% bovine serum albumin diluted in PBS) for 1 h at room temperature under constant rotation for pre-clearing. 1 μg of pre-cleared fragmented DNA in 50 μl TE Buffer was denatured for 5 min at 95°C to obtain single-stranded DNA (ssDNA) and immunoprecipitated overnight at 4°C with 2 μl of anti-8-oxoG monoclonal antibodies or with IgM (Mouse IgM Isotype Control-11E10, ThermoFisher Scientific) or without antibody (No Ab) in a final volume of 250 μl IP buffer (110 mM NaH_2_PO_4_, 110 mM Na_2_HPO ph 7.4, 0.15 M NaCl, 0.05% Triton X-100, 100 mM Tris–HCl pH 8.0, 0.5 M EDTA pH 8.0) under constant rotation. The next day, the immuno-precipitated complex was incubated with 50 μl of pre-saturated Dynabeads Protein G (ThermoFisher Scientific) for 3 h at 4 °C, under constant rotation, and washed three times with 1 ml Washing buffer (110 mM NaH_2_PO_4_, 110mM Na_2_HPO_4_ pH 7.4, 0.15 M NaCl, 0.05% Triton X-100). The bead–antibody–DNA complexes were then disrupted by incubation in 100 μl Lysis buffer (50 mM Tris–HCl pH 8, 10 mM EDTA pH 8, 1% SDS, 0.5 mg/ml Proteinase K) for 4 h at 37°C, and 1 h at 52°C, following the addition of 50 μl Lysis buffer. The immunoprecipitated ssDNA was purified by using MinElute PCR Purification kit (QIAGEN-28004) in a final elution volume of 17 μl in EB buffer (provided in the kit). Conversion of ssDNA to double-stranded DNA (dsDNA) was done using DNA Polymerase I, Large (Klenow) Fragment (Cat. No. M0210S, NEB). Library preparation was performed using the NEBNext® Ultra II DNA Library Kit for Illumina (E7645S) as per manufacturer’s instructions using 25 ng of immunoprecipitated or Input DNA; the only modification was the use of KAPA HiFi Polymerase (Roche) for final library amplification. Prior to sequencing, libraries were quantified using Qubit (Invitrogen) and quality-controlled using Tape Station 4150 (Agilent Technologies). All libraries were multiplexed and run on an Illumina NextSeq 500 as a paired-end run of 150 nt (outsourced to MedGenome Labs Ltd., Bengaluru, India). Note that all of the steps of the OxiDIP-seq protocol, including the washes of the immunocomplexes, were carried out in low-light conditions. Furthermore, 50 μM *N-tert*-butyl-α-phenylnitrone (stock solution: 28 mM in H_2_O; B7263, Sigma) was added to all buffers of the Dneasy Blood and Tissue kit, and IP and washing buffers to preserve the oxidized DNA (Lu et al., 2004).

### Analysis of OxiDIP-seq data

Quality control of the demultiplexed fastq files obtained from Illumina sequencing was performed using FastQC (version 0.11.5), followed by adapter removal and trimming using Trim Galore! Version 0.5.0 (Krueger et al., 2023) with default parameters. Alignments were performed with BWA-MEM (Li & Durbin, 2010) to the *P. falciparum* 3D7 reference genome downloaded from PlasmoDB (https://plasmodb.org, v3, release 68) using default parameters; mapping statistics of the OxiDIP-seq fastq files are provided in **Supplementary Table 1**. SAMtools (Li et al., 2009) and BEDtools (Quinlan & Hall, 2010) were used for filtering steps and file format conversion. Deduplicated sorted bam files filtered for quality (Q > 30) using SAMtools (Li et al., 2009) served as a starting point to identify enriched peaks in immunoprecipitated DNA relative to the gDNA input or IgM/No Ab controls. First, correlation of the bam files of the different stages and replicates was calculated using BEDtools and deepTools (Ramírez et al., 2014). Second, for each timepoint, 8-oxoG-enriched peaks were identified from uniquely mapped reads, after removing duplicates, using the MACS2 software (Y. Zhang et al., 2008) after controlling for a false discovery rate (q-value) of 0.01; three replicates for each timepoint were considered as “treatment” and compared to two input DNA replicates to generate narrowpeaks.bed and a fold-enrichment bedgraph (FE.bedGraph) for every timepoint as previously described (https://github.com/taoliu/MACS/wiki/Build-Signal-Track). The FE.bedGraph was further converted to bigwig and correlation across timepoints was determined using deepTools. The Circos plot was generated using FE.bedGraph files via the Galaxy platform (Afgan et al., 2018). Motif enrichment analysis was performed using the HOMER (Hypergeometric Optimization of Motif EnRichment) software suite (v5.0) (Heinz et al., 2010). Peaks were annotated to genomic features based on overlap with GFF-defined regions in the *P. falciparum* genome (v68) using PeakMatcher (Nowling et al., 2020); a ±20 bp flanking window was applied to accommodate boundary uncertainty and minor positional variation in peak calling. For peaks that overlapped with multiple features, only the maximum overlap was considered. Heatmaps were plotted using R (version 4.4.3).

### RNA-seq library generation, sequencing and analysis

Total RNA extracted from synchronised early ring stage parasites (*i.e.,* 10 hpi) was poly(A) enriched using the NEBNext^®^ Poly(A) mRNA Magnetic Isolation Module (Cat# E7490S). Strand-specific RNA-seq libraries were prepared using the Collibri™ Stranded RNA Library Prep Kit for Illumina™ Systems (Cat# A39116024; ThermoFisher Scientific) according to the manufacturer’s instructions; the only modification was the use of KAPA HiFi Polymerase for final library amplification. Libraries were multiplexed and paired-end sequencing (150 bp) was performed by MedGenome Labs Ltd. using Illumina NextSeq 550, and raw sequencing data were obtained in FASTQ format. Post-quality control using FASTQC (https://www.bioinformatics.babraham.ac.uk/projects/fastqc/), adapter sequences and low-quality bases were removed using Trim Galore! v0.6.4_dev. High-quality reads were aligned to the *P. falciparum* 3D7 reference genome using STAR v2.7.3a (Dobin et al., 2013) with default parameters. SAM files were converted to BAM format using Samtools v1.10 and only reads with a mapping quality score (MAPQ) ≥30 were retained for downstream analysis. Gene-level read counts were generated using HTSeq v0.11.1 (Anders et al., 2015) and gene expression levels were normalized as reads per kilobase of transcript per million mapped reads (RPKM) using an R script.

### Correlation of 8-oxoG levels to RNA-seq, MNase-seq and histone PTM ChIP-seq data

To determine the epigenetic role of 8-oxoG in regulating *P. falciparum* gene expression, the genome-wide distribution profile of 8-oxoG derived from OxiDIP-seq was compared to steady state transcript levels using the in-house-generated RNA-seq data for 10 hpi as well as publicly available 3D7 datasets (Toenhake et al., 2018) for all the time points (10, 20, 30, 40 hpi); the fastq files of Toenhake et al. were downloaded from NCBI SRA (Accession ID PRJNA408158) and processed as described above. First, for each timepoint, genes were classified into two groups based on the presence or absence of 8-oxoG, and gene expression RPKM levels compared between the two groups and visualized in R using ggplot2. Statistical significance between the groups was assessed using the non-parametric Wilcoxon rank-sum test (Wilcoxon, 1945). Next, to compare temporal patterns of 8-oxoG enrichment and gene expression, 8-oxoG FE and RPKM values were log-transformed and normalized using row-wise Z-scores across the four developmental stages. Heatmaps were generated using ggplot2 (Wickham, H. 2016). Hierarchical clustering was performed on the 8-oxoG enrichment matrix, and the resulting gene order was retained for the corresponding expression heatmap to facilitate direct comparison between oxidative DNA modification and transcriptional dynamics during the IDC. Gene Ontology (GO) analysis for genes containing 8-oxoG-enriched peaks was performed using PlasmoDB.

Previously published MNase-seq data (NCBI SRA Accession ID PRJNA133173; Kensche et al., 2014) which had been pre-processed to generate genome-wide nucleosome occupancy profiles by NucMap (Kensche et al., 2016; Zhao et al., 2019) were downloaded as bed files. Correlation of OxiDIP-seq FE.bedgraphs with nucleosome occupancy was plotted using deepTools. Publicly available datasets were also used to compare the 8-oxoG distribution with known patterns of histone PTM and histone variant distribution across the *P. falciparum IDC.* H3K36me2, H3K36me3, and H4K20me3 datasets were retrieved from Jiang et al. (2013) under NCBI SRA accession PRJNA205237. Additional histone marks including H3K4me2, H3K4me3, H3K9ac, H3K14ac, H3K27ac, H4ac, H3K9me3, and H3K27me3 were accessed from Karmodiya et al. (2015) through NCBI SRA accession PRJNA267591. H2A.Z variant histone data were accessed from Bártfai et al (Bártfai et al., 2010; NCBI SRA Accession ID PRJNA133173). All raw data files were downloaded in FASTQ format and processed through a standard analysis pipeline. Briefly, quality control and trimming were performed using FASTQC and TrimGalore, followed by sequence alignment to the *P. falciparum* 3D7 genome using BWA-MEM. Peak calling was carried out with MACS2 using the appropriate Input control files provided in each study. The resulting bedGraph files were converted to BigWig format, which were then used to generate correlation plots between 8-oxoG and histone PTMs using deepTools.

Lastly, 8-oxoG enriched peaks were compared with predicted G-quadruplex-forming sequences in the *P. falciparum* genome, the genomic coordinates of which were obtained from a published study (Gazanion et al., 2020). The comparison was performed using deepTools2, as described.

## Results

### Confirmation of the presence of 8-oxoG in the *P. falciparum* genome

We first assessed the presence of 8-oxoG in genomic DNA isolated from mixed blood-stage *P. falciparum* parasites of the 3D7 laboratory strain using South-western blotting with commercial anti-8-oxoG antibodies. For comparative analysis, we examined 8-oxoG levels in organisms with varying genomic GC-content: humans (white blood cells (WBCs) and the embryonic kidney cell line HEK293; 40.9% GC (Lander et al., 2001)), *Saccharomyces cerevisiae* (38% GC; Goffeau et al., 1996), and *Escherichia coli* (50.8% GC; Blattner et al., 1997). Despite having the lowest GC-content amongst the organisms analyzed, *P. falciparum* genomic DNA exhibited the highest relative 8-oxoG signal intensity, indicating a substantial accumulation of oxidative guanine lesions in the parasite genome (**Figure 1A and 1B**). Thereafter, to examine stage-specific dynamics of 8-oxoG, genomic DNA was isolated from tightly synchronized ring (10-12 hours post-invasion (hpi), trophozoite (28-30 hpi), and schizont (40-42 hpi) stage parasites and analyzed across a DNA concentration gradient. Both qualitative assessment and quantitative measurements consistently revealed a stage-dependent increase of this oxidative mark as the parasite matures within erythrocytes, *i.e.*, genomic 8-oxoG were lowest in rings, higher in trophozoites, and peaked in schizonts (**Figure 1C and 1D**).

**Figure 1:**
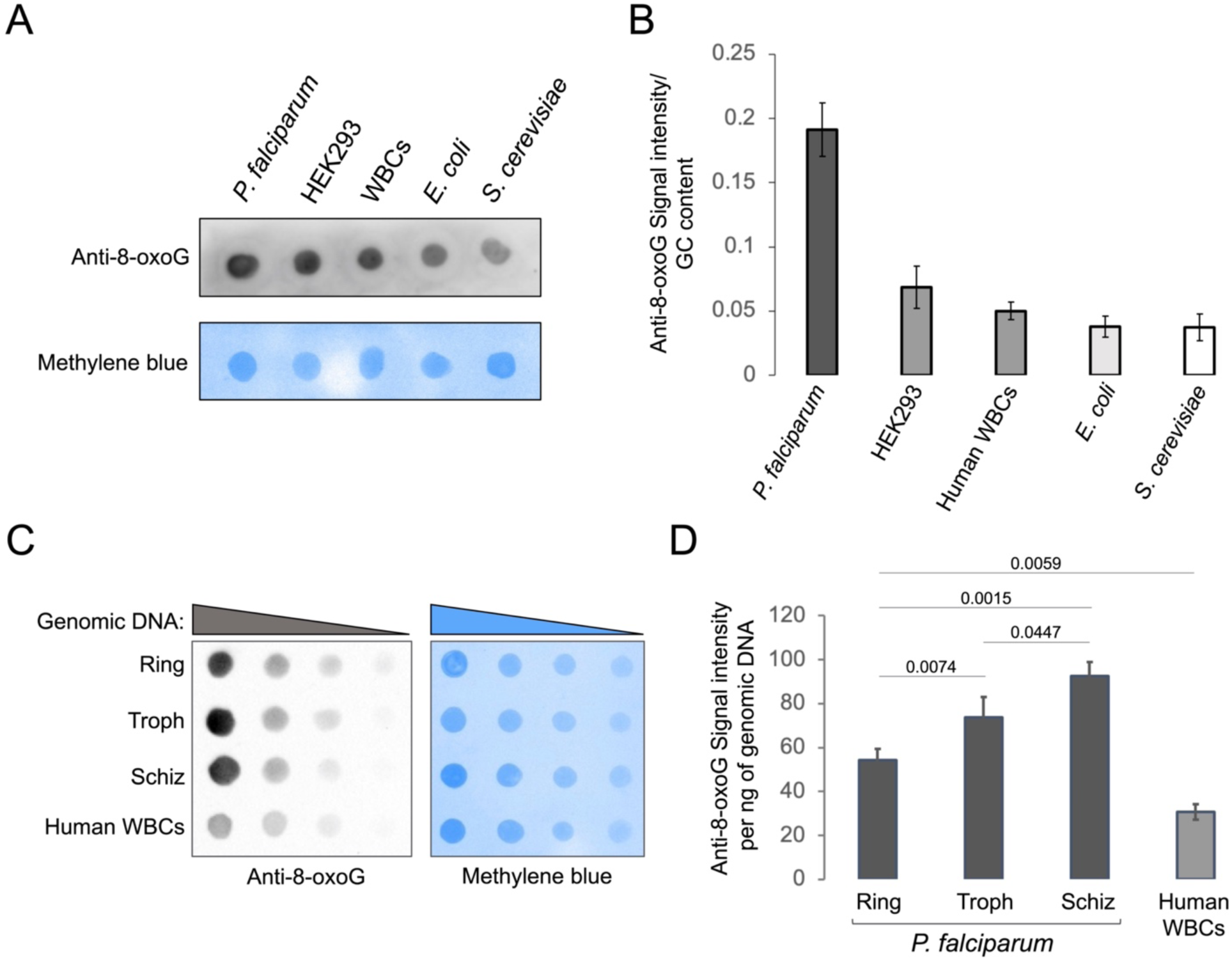
8-oxoG quantification in *P. falciparum* gDNA by South-western blotting. **(A)** Cross-species detection of 8-oxoG in genomic DNA. ***Upper panel:*** Genomic DNA (200 ng) isolated from *P. falciparum*, HEK293 cells, human WBCs, *E. coli*, and *S. cerevisiae* was spotted onto a positively charged nylon membrane and probed with anti-8-oxoG antibodies. ***Lower panel:*** An identical membrane was stained with 0.1% methylene blue to assess DNA loading. **(B)** Quantitative analysis of 8-oxoG signal intensity. Signal intensities from anti-8-oxoG blotting and methylene blue staining were quantified using ImageJ. Normalized signal intensities per ng of genomic DNA and GC content are presented as mean ± SEM (n = 3). **(C)** Stage-specific detection of 8-oxoG in *P. falciparum*. ***Left panel:*** Genomic DNA isolated from synchronized *P. falciparum* 3D7 ring, trophozoite, and schizont stages was spotted in a concentration gradient (400–50 ng) onto a positively charged nylon membrane and probed with anti-8-oxoG antibodies. Human WBC genomic DNA served as a positive control. ***Right panel:*** The corresponding methylene blue-stained blot, showing comparable DNA loading. **(D)** Quantification of stage-specific 8-oxoG levels. Signal intensities were quantified as described in (B), and normalized values are presented as mean ± SEM (n = 5). Statistical significance was determined using Student’s t-test.

### OxiDIP-seq reproducibly enriches 8-oxoG-containing DNA fragments from AT-rich *P. falciparum* genomic DNA

Next, to investigate the genome-wide distribution of 8-oxoG during the *P. falciparum* IDC, a modified version of the OxiDIP-seq method, originally described by the groups of Majello and Amente (Amente et al., 2019a; Gorini et al., 2022) was employed (**Figure 2A**). Because the *P. falciparum* genome is highly AT-rich (∼80.6%), we had to substantially optimize the method (**Supplementary Figure S1**). First, genomic DNA fragmentation was optimized to achieve a size range suitable for downstream sequencing. Two sonication approaches, one using the Covaris S2 focused-ultrasonicator and the other using a bath ultrasonicator, were tested, and both produced fragments within the desired size range of ∼200–600 bp (**Supplementary Figure S2A; *upper panel***). Further, to check the impact of shearing on 8-oxoG levels, *P. falciparum* genomic DNA prepared using two different kits, QIAamp Mini and MagAttract HMW DNA Kits, were tested in South-western dot blotting assays using anti-8-oxoG antibodies; human WBC genomic DNA was used as a control. Remarkably, across all conditions, a consistent reduction in the signal of 8-oxoG was observed post-shearing, with the bath ultrasonicator exhibiting a reduced impact on the 8-oxoG signal as compared to the Covaris instrument (**Supplementary Figure S2A; *lower panel***). Therefore, the bath ultrasonicator was used for all subsequent experiments and a pilot OxiDip-seq run was performed. As shown in **Supplementary Figure S2B**, anti-8-oxoG-immunoprecipitated samples showed a substantially higher 8-oxoG signal compared to the IgM mouse isotype control as well as input DNA, indicating that 8-oxoG–modified DNA fragments can be specifically and efficiently captured using our optimised OxiDIP approach, even from the AT-rich genomic DNA of *P. falciparum*.

**Figure 2:**
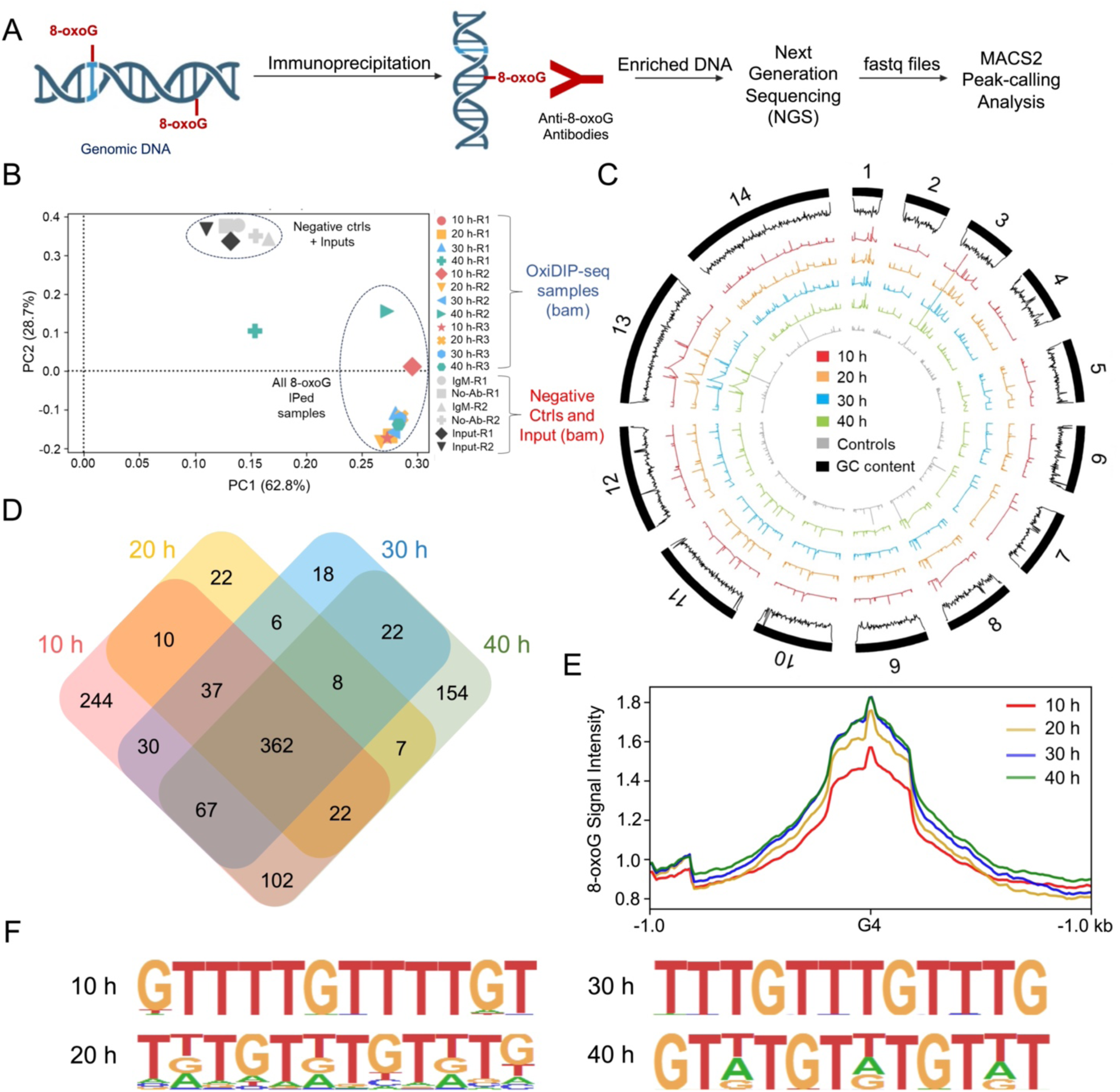
OxiDIP-enriched 8-oxoG genomic loci in *P. falciparum* are GT-rich and correlate with PQSs. **(A)** Schematic overview of the OxiDIP-seq workflow used for genome-wide mapping of 8-oxoG, including genomic DNA isolation, immunoprecipitation with anti-8-oxoG antibodies, library preparation, and sequencing. **(B)** PCA plot of bam alignment files derived from Illumina sequencing data of the OxiDIP-seq biological replicates. The eigenvalues of the top two principal components PC1 (62.8% of variance explained) and PC2 (28.7% of variance explained) are shown and meaningful clustering of samples is indicated. [10 h = early ring, 20 h = late ring, 30 h = trophozoite, 40 h = schizont, ctrls= controls, R1, R2, R3 = biological replicates 1, 2 and 3, respectively]. **(C)** Circos plot showing the distribution of 8-oxoG enrichment across all 14 chromosomes of *P. falciparum*. The outermost thick black segments represent the chromosomal ideograms, with chromosome numbers indicated. The adjacent thin black line denotes GC content across each chromosome. Inner concentric tracks represent MACS2-derived 8-oxoG fold-enrichment profiles at different IDC stages: 10 h, 20 h, 30 h and 40 h (red, orange, blue and green tracks, respectively). The innermost grey track corresponds to the negative control. **(D)** A Venn diagram was used to illustrate the number of shared 8-oxoG-enriched peaks across the 10 h, 20 h, 30 h and 40 h samples. **(E)** Association of 8-oxoG peaks with PQSs, whose coordinates and G4Hunter scores were downloaded from a previously published study (Gazanion et al., 2020). **(F)** Motif enrichment analysis of 8-oxoG–associated regions identified by MACS2 was performed using HOMER. The top-scoring *de novo* motifs for each developmental stage are shown.

Therefore, OxiDIP-seq was performed using genomic DNA isolated from four IDC stages, each with three biological replicates: early rings (10–12 hpi), late rings (20–22 hpi), trophozoites (30–32 hpi) and schizonts (40–42 hpi), with the inclusion of two “no-antibody” controls and two IgM mouse isotype controls. Quality control analysis of the mapped OxiDIP-seq files using principal component analysis (PCA; **Figure 2B**) and Pearson’s Correlation Coefficient (PCC; **Supplementary Figure S3A**) revealed strong clustering and high correlation among all OxiDIP-seq samples, irrespective of parasite stage, indicating robust reproducibility across replicates. In contrast, the negative controls, including input and isotype controls, formed a distinct cluster (**Figure 2B and Supplementary Figure S3A**). Of note, one sample (40 h Replicate1) displayed inconsistent clustering behaviour and was identified as an outlier and therefore excluded from subsequent analyses.

### Genome-wide distribution analysis of 8-oxoG peaks reveals a strong correlation with putative G-quadruplex-forming sequences in the *P. falciparum* genome

To identify 8-oxoG-enriched genomic regions at each timepoint of the *P. falciparum* asexual lifecycle, MACS2 peak-calling analysis of the OxiDIP-seq samples was performed relative to input DNA; parallelly, the control samples were also analysed. Based on PCA and PCC analyses of the MACS2-derived fold-enrichment profiles, a clear separation of OxiDIP-enriched samples from control samples was observed (**Supplementary Figure S3B & S3C**), which was further confirmed by visualizing the profiles using Circos plots (**Figure 2C**). We then calculated the number of 8-oxoG-enriched peaks in the 10 hpi, 20 hpi, 30 hpi, and 40 hpi samples using a cutoff of log_2_(fold change) ≥ 1 and false discovery rate < 0.05, and identified 1574, 866, 1005 and 1381 peaks, respectively (**Supplementary Table S2**). An analysis of shared peaks across all time points indicated that approximately 75% of the peaks are present in two or more stages, with 244, 22, 18 and 154 peaks unique to the 10 hpi, 20 hpi, 30 hpi, and 40 hpi samples, respectively (**Figure 2D**). This suggested that the 8-oxoG DNA base modification remains relatively conserved across the various *P. falciparum* blood stages and exhibits low dynamicity.

Next, to determine whether 8-oxoG is preferentially deposited at G-rich sequences in the *P. falciparum* genome, we performed motif enrichment analysis of the MACS2-derived peaks using HOMER. Both “known” (**Supplementary Figure S4**) and *de novo* motifs (**Figure 2E**) were identified, the latter being particularly valuable given the limited number of characterized transcription factors in *P. falciparum*. The “known” results revealed a G-rich motif conserved in three out of four stages, which corresponds to the ZML2 binding site; ZML2 is a transcription factor in *A. thaliana* but its homolog has not been identified in *P. falciparum* (Shikata et al., 2004). At 30 hpi, a different “known” motif was identified, which corresponds to the Forkhead Box L1 (FOXL1) transcription factor recognition site in humans (Fukuda et al., 2003). In contrast, the *de novo* motifs were GT-rich (**Figure 2E**) and varied across the four time points. Given this observation and considering that studies in mammalian cells have shown that 8-oxoG–containing peaks are highly enriched at G quadruplex-forming sequences (Fleming & Burrows, 2020; Kuznetsova et al., 2020; Zhou et al., 2013), we measured 8-oxoG signal intensity at putative G quadruplex-forming sequences (PQS) of the *P. falciparum* genome using a published dataset of 1763 PQSs at a threshold of 1.2 (Gazanion et al., 2020). As shown in **Figure 2F**, we found that, for all four IDC timepoints, the 8-oxoG signal accumulated around PQSs, indicating that 8-oxoG deposition at PQSs may be an ancient phenomenon.

### 8-oxoG modifications occur predominantly in coding regions of *P. falciparum* protein-coding genes and exhibit a marked enrichment at gene 3’ ends

To study the genomic features associated with 8-oxoG during the *P. falciparum* IDC, peak annotation was performed, which revealed that a majority of the 8-oxoG-containing peaks map to exonic regions followed by intergenic regions and 5’/3’ UTRs (**Figure 3A**). A deeper analysis of the protein-coding genes marked with 8-oxoG at 10 hpi, 20 hpi, 30 hpi, and 40 hpi, revealed that 931, 520, 601, and 787 genes, respectively, are marked with 8-oxoG within exons, introns, and/or 5’/3’ UTRs, with multigene families comprising approximately 15% of the total 8-oxoG-marked genes (**Figure 3B**). An analysis of the shared genes across all time points indicated that approximately 35-45% of the modified genes have 8-oxoG peaks in all IDC stages, with 35-45% having 8-oxoG peaks in 3 of 4 stages, and 10-20% in 2 of 4 stages (**Figure 3C**). Of note, the ∼300 core genes (after removing multigene families) associated with 8-oxoG across all timepoints include all three members of the pre-slicing U2AF complex, three calcium-dependent protein kinases (*PF3D7_0317200, PF3D7_0415300*, and *PF3D7_1356900*), eight DNA repair pathway genes (*PF3D7_0305700, PF3D7_0317700, PF3D7_0408500, PF3D7_0416300, PF3D7_0623900, PF3D7_0803400, PF3D7_1244200* and *PF3D7_1332600*) and 21 genes encoding for mRNA binding/regulatory proteins (**Supplementary Table S3**).

**Figure 3:**
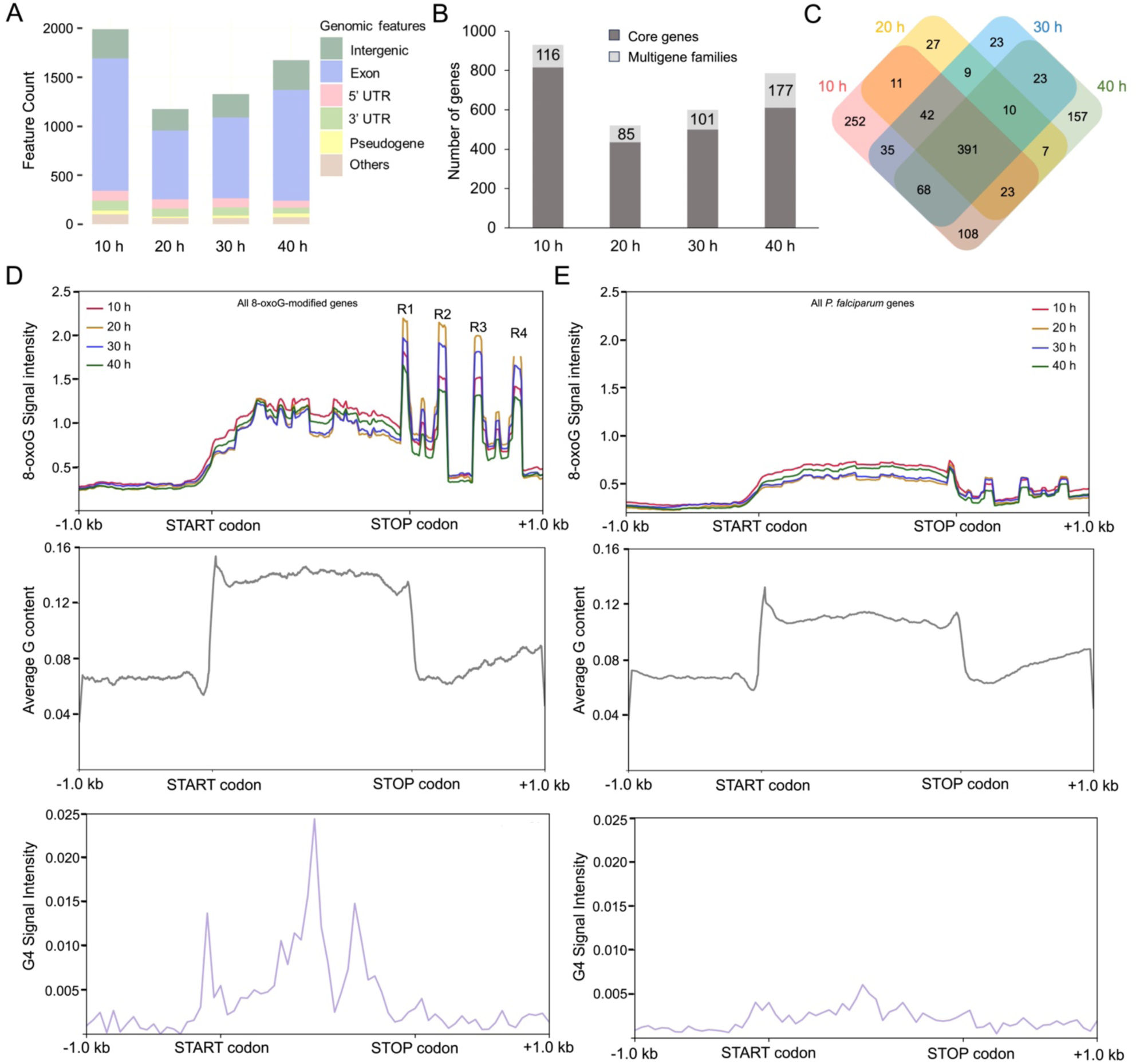
8-oxoG modifications occur predominantly in exonic regions of protein-coding genes and show low dynamicity. **(A)** The number of 8-oxoG-enriched peaks overlapping with different genomic features like intergenic regions, exons, introns, 5’UTRs, 3’UTRs and pseudogenes is shown for the indicated time points. **(B)** The bar graph shows the total number of core genes and genes belonging to multigene families with annotated 8-oxoG peaks at each developmental stage. **(C)** A Venn diagram was used to depict the number of shared genes across the different IDC stages. **(D**) ***Upper panel:*** Metagene profiles showing the average distribution of 8-oxoG signal across all modified genes, from 1 kb upstream of the START codon (ATG) to 1 kb downstream of the STOP codon (TAA). R1, R2, R3 and R4 indicate highly enriched peaks around the STOP codon. ***Middle panel:*** Average G-content (%) across the same genomic regions. ***Lower panel:*** Average density of PQSs obtained from Gazanion et al., 2020 across the same genomics regions. **(E)** Same as (D), but for all *P. falciparum* protein-coding genes.

Subsequent calculation of the average 8-oxoG density profile across all modified genes, while confirming higher enrichment within coding regions (*i.e.,* between the START and STOP codons), revealed a pronounced increase in 8-oxoG signal close to the STOP codon, which manifested in four distinct regions: the first enriched region R1 is located upstream of the STOP codon and the remaining three regions R2 to R4 are present downstream of the STOP codon (**Figure 3D, top panel**). To further investigate this pattern, we zoomed into the region between the STOP codon and transcription end site (TES), *i.e.,* the 3′ UTR, and observed that only R2 falls within the 3’UTR, while R3 and R4 are present downstream of the TES (**Supplementary Figure S5**). To assess whether the observed genic 8-oxoG enrichment pattern is driven by nucleotide composition, we analyzed the average G content across 8-oxoG-modified genes and found it to be relatively higher within the coding sequence as compared to the region downstream of the STOP codon (**Figure 3D, *middle panel***), in line with published studies (Gardner et al., 2002b). Similarly, PQS density was also higher within the coding sequence of 8-oxoG-modified genes relative to their 3’ flanks (**Figure 3D, *lower panel***). This was in contrast to lower 8-oxoG signal density, G content and PQS density, on average, across all *P. falciparum* protein-coding genes (**Figure 3E**). Taken together, we conclude that 8-oxoG is actively targeted to select *P. falciparum* genomic regions rather than being a mere stochastic product of nucleotide composition.

### 8-oxoG enrichment correlates with increased steady-state transcript levels and stage-specific gene expression

Previous studies in mammalian systems have correlated 8-oxoG accumulation within G-quadruplex sequences with gene activation (Fleming, Zhu, et al., 2017; Gorini et al., 2023). Therefore, to establish the gene regulatory potential of 8-oxoG modifications in *P. falciparum*, we examined whether the presence of 8-oxoG within the gene body is associated with higher or lower steady state transcript levels during asexual blood stages. For this analysis, we compared 8-oxoG-modified genes to unmodified genes using publicly available transcriptomic datasets for 20 hpi and 40 hpi of the *P. falciparum* 3D7 strain (Toenhake et al., 2018) and found that genes harbouring 8-oxoG peaks exhibited significantly higher steady-state transcript levels when compared to genes lacking these modifications (**Figure 4A**). To further examine the relationship between 8-oxoG enrichment and gene expression, the log₂ fold enrichment (log₂FE) values of 8-oxoG–modified genes were compared across 10, 20, 30 and 40 hpi using z-score-based hierarchical clustering and integrated with their corresponding RPKM-normalized and z-score-transformed RNA-seq data (from Toenhake et al., 2018). The combined analysis revealed the following: (1) 8-oxoG enrichment follows a cyclical pattern during the IDC, based on which the 8-oxoG-modified genes can be divided into four clusters, cluster 1 corresponding to maximal 8-oxoG enrichment at 40 hpi, cluster 2 at 10 hpi, cluster 3 at 20 hpi and cluster 4 at 30 hpi (**Figure 4A**). (2) A majority of the 8-oxoG-modified genes exhibit maximal steady-state transcript levels in the stage when log₂FE values of 8-oxoG are the highest (**Figure 4B**) suggesting a transcriptional activation role for 8-oxoG during the IDC.

**Figure 4:**
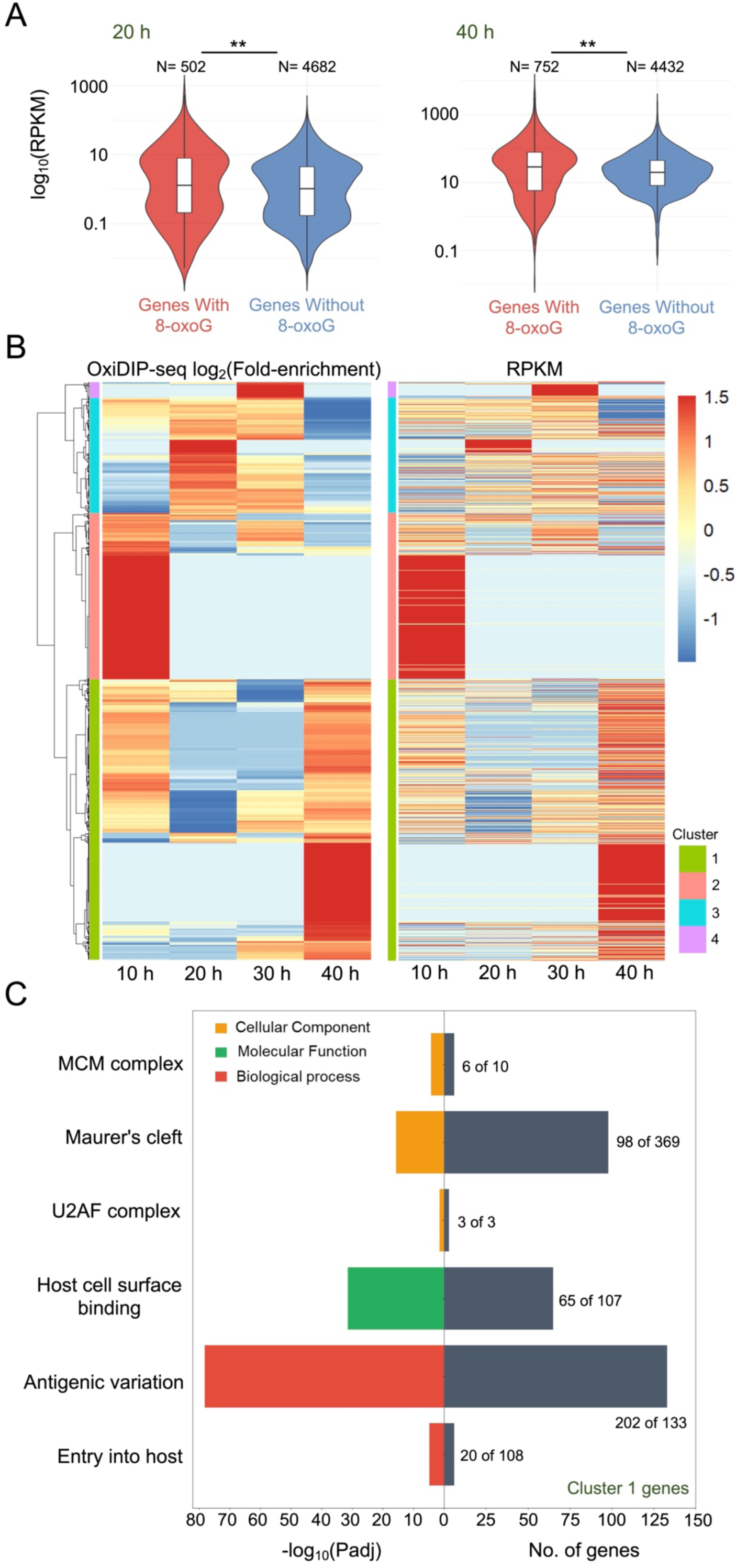
The presence of 8-oxoG in genic regions is associated with elevated steady-state transcript levels. **(A)** Steady state transcript levels of genes with or without 8-oxoG modifications were compared using publicly available RNA-seq datasets from Toenhake et al., 2018, for 20 h and 40 h. The indicated p-values were calculated from a Wilcoxon test with ** corresponding to p-value <0.01. N indicates the total number of genes in each category. **(B)** Heatmaps showing the hierarchical clustering of modified genes based on log-transformed and z-normalised 8-oxoG fold enrichment across four IDC stages (***left panel***) and the corresponding log-transformed and z-normalised steady-state transcript levels for the same genes using the 10 h, 20 h, 30 h and 40 h RNA-seq datasets of Toenhake et al., 2018 (***right panel***). Each row represents an individual gene and columns correspond to the indicated time points. The color scale indicates normalized signal intensity (z-score), ranging from low (blue) to high (red). Genes are grouped into distinct clusters, indicated by colored bars on the right. **(C)** GO enrichment analysis of genes from cluster 1 of part B was performed using PlasmoDB. Significant GO terms (Benjamini-adjusted p-value or Padj ≤ 0.05) related to Molecular Function (green), Cellular Component (yellow), and Biological Process (grey) were plotted with the number of genes enriched for each GO term relative to background indicated on the right y-axis, while the -log_10_(Padj) of each GO term is shown on the left y-axis.

Next, to determine the functional categories of the genes belonging to each cluster, gene ontology (GO) overrepresentation analysis was performed using PlasmoDB (https://plasmodb.org). Only Cluster 1, which represents the largest gene set, showed overrepresented GO terms for biological processes associated with antigenic variation, and pre-replicative complex assembly, molecular functions such as host cell surface binding, and cellular components such as U2AF complex, MCM complex and Maurer’s cleft (**Figure 4C**). Overall, these results demonstrate that 8-oxoG modifications are not randomly distributed across the *P. falciparum* genome, but are preferentially localized to actively transcribed genes that belong to important virulence pathways. Of note, when multigene families such as *var* (**Supplementary Figure S6A)** and *rifin* (**Supplementary Figure S6B)** were specifically compared at the 8-oxoG log₂FE and RPKM levels, we did not observe a clear association between 8-oxoG genic occupancy and steady state transcript levels.

### 8-oxoG-marked genomic loci preferentially associate with active and poised histone PTMs

Having established the transcriptional activating potential for 8-oxoG, we next asked if there is a correlation between 8-oxoG enrichment and chromatin organization, for which 8-oxoG profiles were first compared with nucleosome occupancy profiles extracted from MNase-seq data for 20 and 40 hpi (Kensche et al., 2016; Zhao et al., 2019). This comparison revealed that 8-oxoG-marked genomic loci do not occlude nucleosomes (**Figure 5A**), which was further confirmed by analysing 8-oxoG signals within nucleosome-bound regions of the genome relative to linker DNA (**Figure 5B**). In fact, 8-oxoG and unmodified G showed a similar pattern of nucleosome positioning suggesting that the presence of 8-oxoG does not substantially alter global nucleosome organization or chromatin accessibility within the parasite genome.

**Figure 5:**
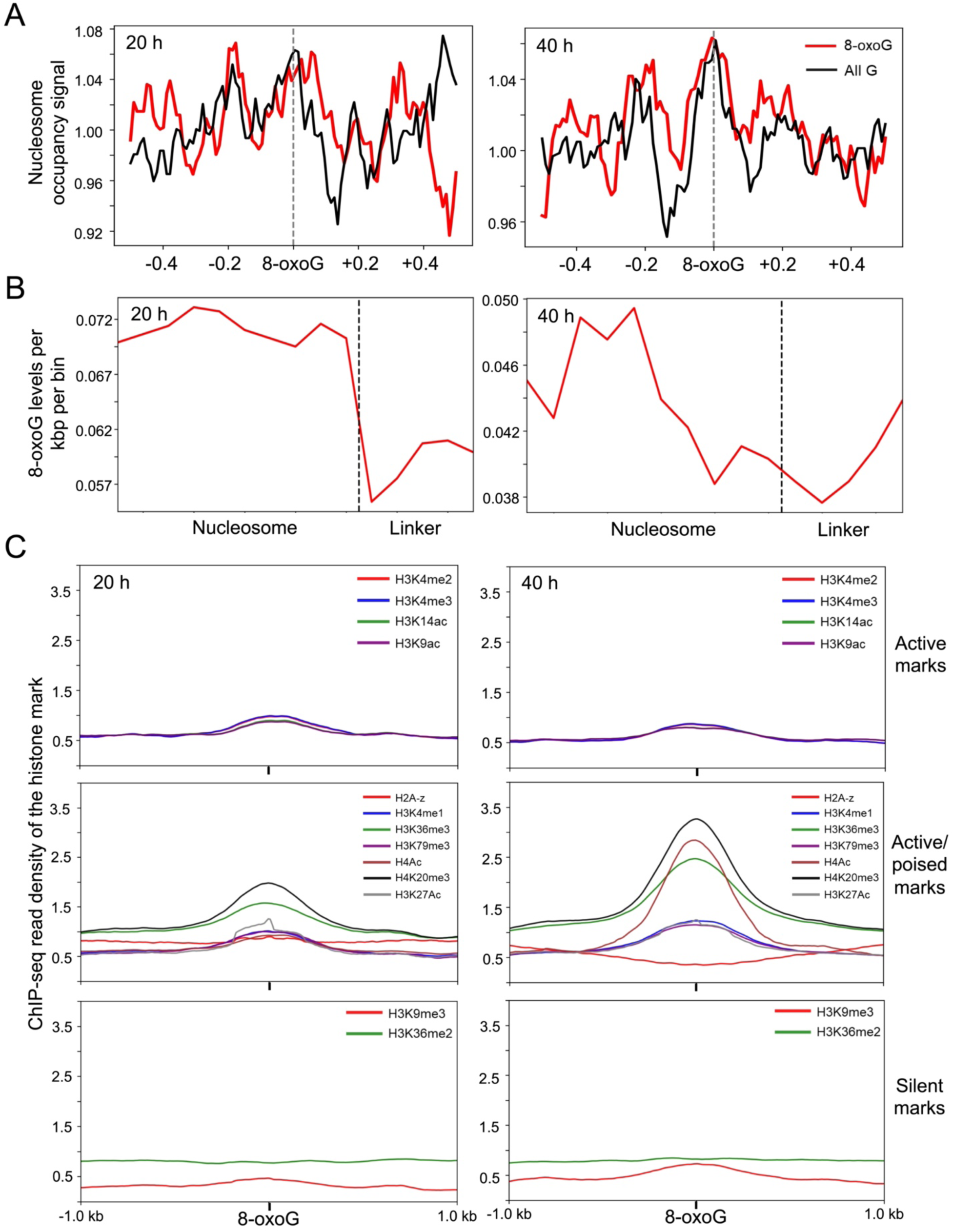
8-oxoG-containing peaks in the *P. falciparum* genome show a positive correlation with active/poised histone PTMs. (A) Average profiles of nucleosome occupancy at 8-oxoG-enriched loci for 20 h (*left panel)* and 40 h (*right panel*) samples. The X-axis represents a guanosine base anchored at “0” and its +/- 500 bp flanking regions. The Y-axis represents the nucleosome occupancy signal obtained from Kensche et al., 2016. The red line is centred at 8-oxoG loci while the black line is centred at an equal number of guanosine sites chosen at random. **(B)** Average 8-oxoG profile across nucleosome and linker regions at 20 h (*left panel)* and 40 h (*right panel*). The X-axis represents the position of the nucleosome (divided into 10 bins) and linker (divided into 4 bins). The Y-axis represents 8-oxoG levels per kbp per bin. **(C)** DeepTools plotprofile was used to compare the read density distribution of the indicated active, active/poised and silencing histone PTMs for all 8-oxoG genomic positions located within the exons of modified genes at 20 h (*left panel)* or 40 h (*right panel*). Profiles represent normalized signal intensity of the histone PTM across the indicated regions.

Consequently, we asked whether 8-oxoG-enriched genomic regions are associated with specific histone PTMs, particularly those linked to transcriptional activation. Using histone PTM datasets from two published studies (Bártfai et al., 2010a; Jiang et al., 2013; Karmodiya et al., 2015), we analysed the correlation of 8-oxoG to activating PTMs (H3K4me2, H3K4me3, H3K9ac and H3K14ac), active/poised PTMs (H2A.z, H3K4me1, H3K36me3, H3K79me3, H4Ac, H4K20me3 and H3K27Ac), and silencing PTMs (H3K9me3 and H3K36me2) at 20 and 40 hpi for 8-oxoG-modified genes; genes that belong to the *var, rifin* and *stevor* multigene families were excluded from this analysis because these loci are subject to specialized clonally variant epigenetic regulation which may obscure broader genome-wide associations between 8-oxoG enrichment and histone PTM landscapes. As shown in **Figure 5C**, 8-oxoG-modified regions of protein-coding genes show a preferential association with active or poised chromatin states in *P. falciparum* at 40 hpi and to a lesser extent at 20 hpi. Interestingly, when we focused on genes that belong to cluster 1 from Figure 4B, *i.e.,* genes with the highest 8-oxoG levels at 40 hpi, we observed that active/poised histone PTMs are most correlated to 8-oxoG at 40 hpi and not at 20 hpi **(Supplementary Figure S7)**. On the whole, these findings suggest that 8-oxoG preferentially accumulates within transcriptionally permissive chromatin environments and exhibits a dynamic, stage-specific association with active and poised histone PTMs.

## Discussion

Oxidative DNA modifications have long been regarded as unavoidable by-products of ROS and have been investigated primarily for their mutagenic potential and contribution to genome instability (Kino et al., 2017). However, growing evidence from higher eukaryotes has demonstrated that specific oxidative DNA lesions, particularly 8-oxoG, can also participate in the regulation of gene expression through interactions with chromatin architecture, DNA repair proteins, transcription factors, and epigenetic regulatory pathways (Amente et al., 2010; Ba & Boldogh, 2018b; Gorini et al., 2020; Hahm et al., 2022). Despite these advances, whether oxidative DNA modifications contribute to epigenetic regulation in evolutionarily divergent eukaryotes such as *P. falciparum*, which tightly coordinates its gene expression program during the 48-hour IDC (Bozdech et al., 2003), remains largely unexplored.

To fill this knowledge gap, we first quantified genomic 8-oxoG levels across the *P. falciparum* IDC and compared them with representative prokaryotic and eukaryotic model organisms. Surprisingly, despite possessing the most AT-rich genome amongst the organisms considered, *P. falciparum* showed markedly elevated levels of genomic 8-oxoG, indicating that susceptibility to oxidation-driven DNA base modifications is not determined solely by genomic guanine content. Instead, it may reflect a combination of sustained oxidative stress encountered by the parasite within the RBC and/or limited repair of oxidized DNA lesions. Indeed, as the IDC progresses, the parasite experiences an increasingly oxidative intracellular environment generated by haemoglobin degradation, haem metabolism, mitochondrial respiration, and host-derived ROS (Becker et al., 2004; Francis et al., 1997). Consistent with this, we found that genomic 8-oxoG levels increased progressively throughout the IDC, reaching their highest abundance in schizonts. Moreover, unlike yeast and mammals, which possess an efficient OGG1-mediated BER pathway that rapidly recognizes and removes 8-oxoG in nuclear DNA (David et al., 2007; Hazra et al., 2007), several enzymes of the *P. falciparum* BER machinery localize predominantly to the DNA-containing organelles, apicoplast and mitochondrion, and have not been detected in the nucleus (Haltiwanger et al., 2000; Tiwari et al., 2020, 2024; Verma et al., 2021). Together, these findings suggest that the persistent accumulation of 8-oxoG in the *P. falciparum* genome is more than a simple consequence of oxidative damage and instead raises the possibility that this modified base has been co-opted for regulatory functions during parasite development.

To address this further, we set out to adapt the genome-wide analysis method OxiDIP-seq to *P. falciparum.* Because this technique was originally developed for mammalian genomes with substantially higher GC content (Amente et al., 2019; Gorini et al., 2022), its application to the highly AT-rich *P. falciparum* genome required methodological optimization. Firstly, fragmentation conditions were optimised, wherein bath ultrasonication was found to reduce the genomic 8-oxoG signal to a lesser extent than high-energy focused ultrasonication. Secondly, the chemically labile nature of 8-oxoG (Cadet et al., 2010), necessitated the addition of *N-tert*-butyl-α-phenylnitrone to all the buffers used in the workflow. Thirdly, pre-clearing of genomic DNA strongly reduced background signal in the control IP reactions, possibly by minimizing non-specific antibody interactions and improving enrichment specificity. Consequently, our optimised workflow resulted in a clear separation of OxiDIP-seq libraries from input and isotype IgM controls, together with significant biological reproducibility across replicates. Collectively, we conclude that 8-oxoG profiling using antibodies can be successfully adapted to a genome with low GC content, thereby expanding the epigenomic toolkit available for investigating non-canonical DNA modifications in other AT-rich malaria parasites as well as unrelated pathogens with biased genomic content.

The resultant genome-wide map revealed that 8-oxoG is stably enriched at specific genomic loci throughout the *P. falciparum* IDC, rather than being randomly dispersed as would be expected for oxidative DNA damage. In fact, the GT-rich nature of 8-oxoG-containing motifs in the *P. falciparum* genome and the preferential association of 8-oxoG with PQSs is similar to reports in mammalian systems, where 8-oxoG accumulation is observed within G-rich regions of regulatory DNA elements that are capable of forming G-quadruplex structures (Fleming, Zhu, et al., 2017; Fleming & Burrows, 2021; Kuznetsova et al., 2020). Oxidation of guanine within these regions has been shown to alter G-quadruplex stability, recruit OGG1, and promote local chromatin remodelling as well as transcriptional regulation (Pan, Vlahopoulos, et al., 2023; Visnes et al., 2018). Extrapolating to *P. falciparum,* we speculate that oxidative DNA modifications may contribute to the regulation of parasite genomic architecture and that the G quadruplex-OGG1 relationship may be evolutionarily conserved.

Another noteworthy observation is the strong enrichment of 8-oxoG loci around the 3′ end of select *P. falciparum* genes. This raises the possibility that localized oxidation near gene termini may influence chromatin architecture and/or transcription termination efficiency in turn impacting transcript levels. Alternatively, elevated 8-oxoG near STOP codons may reflect increased formation of non-canonical DNA structures at the ends of protein-coding sequences due to transcription-associated oxidative processes. Distinguishing between these possibilities will require direct biochemical characterization of oxidized loci, OGG1 and associated chromatin factors. Given that 3′UTRs of mRNA transcripts are increasingly recognized as regulatory hubs controlling mRNA stability, transcript localization, translational efficiency and transcription termination (Mayr, 2017; Tian & Manley, 2017), and in *P. falciparum*, as central mediators of post-transcriptional regulation and developmental timing (Bozdech et al., 2003; Bunnik et al., 2013; Le Roch et al., 2004; Painter et al., 2018; Vembar et al., 2016), it will be important to investigate whether 8-oxoG accumulation at gene termini contributes to the coordination of transcriptional and post-transcriptional regulatory mechanisms in *P. falciparum*. Such a role would place oxidative DNA modifications at a strategic interface between chromatin-based regulation and mRNA fate determination during parasite intra-erythrocytic development.

A major finding of this study is the strong association between genic 8-oxoG enrichment and steady-state transcript levels throughout the *P. falciparum* IDC. Genes marked with 8-oxoG showed significantly higher steady-state transcript levels as compared to unmodified genes, and temporal changes in 8-oxoG occupancy closely mirrored stage-specific expression dynamics, with the clearest association observed at 40 hpi. These observations are consistent with studies in other eukaryotes, where 8-oxoG promotes transcription independently (Obermann et al., 2025) or through OGG1-mediated recruitment of transcription factors and modulation of chromatin accessibility (Amente et al., 2019b; Ba & Boldogh, 2018b; Pan, Vlahopoulos, et al., 2023). Moreover, GO enrichment analysis indicated that 8-oxoG-associated genes are involved in genome maintenance, transcriptional regulation, cell cycle progression, and antigenic variation, suggesting a role for this DNA base modification in coordinating key developmental and adaptive processes. This is in agreement with human studies, where 8-oxoG was shown to mark genes involved in oxidative stress response, cellular differentiation, proliferation and homeostasis (Aguilera-Aguirre et al., 2015; Pan et al., 2016; Pan, Hao, et al., 2023). Lastly, given that the *var* and *rifin* multigene families showed no clear relationship between 8-oxoG enrichment and transcript abundance, we conclude that 8-oxoG contributes selectively to specific *P. falciparum* transcriptional programs rather than functioning as a universal activation mark across the parasite genome.

Thereafter, when 8-oxoG-modified genomic loci were analysed for the presence of different classes of histone PTMs/histone variants, we found preferential co-localization of 8-oxoG with H3K4 methylation, histone acetylation marks and the histone H2A.Z variant, all of which have previously been linked to transcriptional activation and developmental progression in blood-stage *P. falciparum* parasites (Bártfai et al., 2010b; Jiang et al., 2013; Karmodiya et al., 2015). In contrast, nucleosome occupancy was not altered at 8-oxoG-enriched loci, suggesting that the relationship between oxidation and transcription may operate primarily through chromatin state transitions rather than large-scale nucleosome repositioning. These findings are particularly significant given that, unlike mammalian genomes where DNA cytosine methylation (*i.e.,* 5mC) constitutes a major epigenetic regulatory mechanism, *P. falciparum* contains only trace levels of 5mC (Hammam et al., 2020; Ponts et al., 2013) and appears to rely heavily on alternative chromatin-based regulatory pathways such as histone PTMs, histone variants, chromatin remodelers, nuclear organization and non-coding RNAs (Coetzee et al., 2017; Connacher et al., 2021; Cui et al., 2015b; Cui & Miao, 2010; Duffy et al., 2014; Gómez-Díaz et al., 2017). The identification of 8-oxoG as a candidate DNA-based epigenetic mark associated with active chromatin and elevated transcription therefore expands the repertoire of molecular mechanisms that may contribute to epigenetic regulation in malaria parasites. When considered alongside our previous work on a putative 5hmC-like modification that is enriched within highly expressed genes (Hammam et al., 2020) and a gene-activating role for N6-methyladenine during the *P. falciparum* IDC (Seshan et al., under review), our results support an emerging model in which non-canonical DNA modifications act alongside histone-based mechanisms to fine-tune *P. falciparum* developmental gene expression, enabling transcriptional adaptation to the fluctuating redox environment encountered by the parasite during blood-stage infection.

Overall, our study provides the first genome-wide characterization of 8-oxoG in *P. falciparum* and reveals that this oxidative DNA base modification exhibits many of the hallmarks of a regulatory epigenetic feature rather than a purely deleterious DNA lesion, adding to a growing body of 8-oxoG work in organisms other than humans. Although important questions remain regarding the precise molecular mechanisms involved in 8-oxoG-mediated gene regulation in *P. falciparum*, for example, the contribution of the putative parasite OGG1 homolog (encoded by *PF3D7_0917100*), the impact of 8-oxoG on chromatin organisation, and the causal, rather than correlative, relationship between oxidation and transcriptional activation, our work supports a model in which oxidative DNA chemistry forms an integral component of the parasite regulatory landscape. From an evolutionary perspective, such a strategy may be particularly advantageous for an organism that completes its asexual and sexual developmental cycles within the highly oxidative environment of the human erythrocyte, allowing environmental redox cues to be directly integrated into transcriptional decision-making. To this end, future work should combine orthogonal 8-oxoG-mapping approaches and functional perturbation of *P. falciparum* OGG1 homologs/orthologs to establish mechanistic links between oxidative DNA chemistry and gene regulation, particularly in the context of G-quadruplex stability, transcription termination and chromatin accessibility. More broadly, our findings substantiate that oxidative DNA modifications can serve as regulatory information carriers across deeply divergent eukaryotic lineages and raise the possibility that the repurposing of oxidative DNA lesions for gene regulation represents an ancient and evolutionarily conserved mechanism for environmental adaptation.

## Data availability

The OxiDIP-seq FASTQ files generated in this study have been deposited in the NCBI Sequence Read Archive (SRA) under BioProject accession **PRJNA1260506**. The RNA-seq FASTQ files analyzed in this study, which were generated as part of another study, are available in the NCBI Sequence Read Archive (SRA) under BioProject accession **PRJNA1260746**.

## Acknowledgements

D.A. acknowledges PhD support from the Department of Biotechnology, India, under the DBT-JRF scheme. We thank Seshan D., Anitha N. Bavikatte, Jayanth Vegesna and Bhavesh Sharma for data analysis inputs.

## Author Contributions

S.S.V. conceived and designed the experiments; D.A. performed all of the experiments except Illumina sequencing; D.A. and S.S.V. analyzed the data and performed statistical analyses; D.A. and S.S.V. wrote the paper and approved the final manuscript.

## Funding Statement

This work was supported by funding from the Department of Electronics, IT, BT & ST, Government of Karnataka, to IBAB, Bengaluru, the Ramalingaswami Re-entry Fellowship (BT/RLF/Re-entry36/2017) from the Department of Biotechnology, Government of India, to S.S.V. and a SERB-CRG grant (CRG/2020/004801) from the Department of Science and Technology, Government of India, to S.S.V.

## Competing Interests Declaration

The authors declare no competing interests.

## Notes

### Competing Interest Statement

The authors have declared no competing interest.

## References

Afgan, E., Baker, D., Batut, B., van den Beek, M., Bouvier, D., Cech, M., Chilton, J., Clements, D., Coraor, N., Grüning, B. A., Guerler, A., Hillman-Jackson, J., Hiltemann, S., Jalili, V., Rasche, H., Soranzo, N., Goecks, J., Taylor, J., Nekrutenko, A., & Blankenberg, D. (2018). The Galaxy platform for accessible, reproducible and collaborative biomedical analyses: 2018 update. Nucleic Acids Research, 46(W1), W537–W544. 10.1093/nar/gky379

Aguilera-Aguirre, L., Hosoki, K., Bacsi, A., Radák, Z., Sur, S., Hegde, M. L., Tian, B., Saavedra-Molina, A., Brasier, A. R., Ba, X., & Boldogh, I. (2015). Whole transcriptome analysis reveals a role for OGG1-initiated DNA repair signaling in airway remodeling. Free Radical Biology and Medicine, 89, 20–33. 10.1016/j.freeradbiomed.2015.07.007

Allgayer, J., Kitsera, N., Bartelt, S., Epe, B., & Khobta, A. (2016). Widespread transcriptional gene inactivation initiated by a repair intermediate of 8-oxoguanine. Nucleic Acids Research, 44(15), 7267–7280. 10.1093/nar/gkw473

Amente, S., Di Palo, G., Scala, G., Castrignanò, T., Gorini, F., Cocozza, S., Moresano, A., Pucci, P., Ma, B., Stepanov, I., Lania, L., Pelicci, P. G., Dellino, G. I., & Majello, B. (2019a). Genome-wide mapping of 8-oxo-7,8-dihydro-2′-deoxyguanosine reveals accumulation of oxidatively-generated damage at DNA replication origins within transcribed long genes of mammalian cells. Nucleic Acids Research, 47(1), 221–236. 10.1093/nar/gky1152

Amente, S., Di Palo, G., Scala, G., Castrignanò, T., Gorini, F., Cocozza, S., Moresano, A., Pucci, P., Ma, B., Stepanov, I., Lania, L., Pelicci, P. G., Dellino, G. I., & Majello, B. (2019b). Genome-wide mapping of 8-oxo-7,8-dihydro-2′-deoxyguanosine reveals accumulation of oxidatively-generated damage at DNA replication origins within transcribed long genes of mammalian cells. Nucleic Acids Research, 47(1), 221–236. 10.1093/nar/gky1152

Amente, S., Lania Luigi, Avvedimento Enrico Vittorio, & and Majello, B. (2010). DNA oxidation drives Myc mediated transcription. Cell Cycle, 9(15), 3074–3076. 10.4161/cc.9.15.12499

Anders, S., Pyl, P. T., & Huber, W. (2015). HTSeq—A Python framework to work with high-throughput sequencing data. Bioinformatics, 31(2), 166–169. 10.1093/bioinformatics/btu638

Ba, X., Bacsi, A., Luo, J., Aguilera-Aguirre, L., Zeng, X., Radak, Z., Brasier, A. R., & Boldogh, I. (2014). 8-Oxoguanine DNA glycosylase-1 augments proinflammatory gene expression by facilitating the recruitment of site-specific transcription factors. Journal of Immunology, 192(5), 2384–2394. 10.4049/jimmunol.1302472

Ba, X., & Boldogh, I. (2018a). 8-Oxoguanine DNA glycosylase 1: Beyond repair of the oxidatively modified base lesions. Redox Biology, 14, 669–678. 10.1016/j.redox.2017.11.008

Ba, X., & Boldogh, I. (2018b). 8-Oxoguanine DNA glycosylase 1: Beyond repair of the oxidatively modified base lesions. Redox Biology, 14, 669–678. 10.1016/j.redox.2017.11.008

Bártfai, R., Hoeijmakers, W. A. M., Salcedo-Amaya, A. M., Smits, A. H., Janssen-Megens, E., Kaan, A., Treeck, M., Gilberger, T.-W., Françoijs, K.-J., & Stunnenberg, H. G. (2010a). H2A.Z Demarcates Intergenic Regions of the Plasmodium falciparum Epigenome That Are Dynamically Marked by H3K9ac and H3K4me3. PLoS Pathogens, 6(12), e1001223. 10.1371/journal.ppat.1001223

Bártfai, R., Hoeijmakers, W. A. M., Salcedo-Amaya, A. M., Smits, A. H., Janssen-Megens, E., Kaan, A., Treeck, M., Gilberger, T.-W., Françoijs, K.-J., & Stunnenberg, H. G. (2010b). H2A.Z demarcates intergenic regions of the plasmodium falciparum epigenome that are dynamically marked by H3K9a<otherinfo>c and H3K4me3. PLoS Pathogens, 6(12), e1001223. 10.1371/journal.ppat.1001223

Batugedara, G., Lu, X. M., Bunnik, E. M., & Le Roch, K. G. (2017). The Role of Chromatin Structure in Gene Regulation of the Human Malaria Parasite. Trends in Parasitology, 33(5), 364–377. 10.1016/j.pt.2016.12.004

Becker, K., Tilley, L., Vennerstrom, J. L., Roberts, D., Rogerson, S., & Ginsburg, H. (2004). Oxidative stress in malaria parasite-infected erythrocytes: Host–parasite interactions. International Journal for Parasitology, 34(2), 163–189. 10.1016/j.ijpara.2003.09.011

Blattner, F. R., Plunkett, G., Bloch, C. A., Perna, N. T., Burland, V., Riley, M., Collado-Vides, J., Glasner, J. D., Rode, C. K., Mayhew, G. F., Gregor, J., Davis, N. W., Kirkpatrick, H. A., Goeden, M. A., Rose, D. J., Mau, B., & Shao, Y. (1997). The complete genome sequence of Escherichia coli K-12. Science, 277(5331), 1453–1462. 10.1126/science.277.5331.1453

Boiteux, S., & Radicella, J. P. (2000). The Human *OGG1* Gene: Structure, Functions, and Its Implication in the Process of Carcinogenesis. Archives of Biochemistry and Biophysics, 377(1), 1–8. 10.1006/abbi.2000.1773

Boland, M. J., Nazor, K. L., & Loring, J. F. (2014). Epigenetic Regulation of Pluripotency and Differentiation. Circulation Research, 115(2), 311–324. 10.1161/CIRCRESAHA.115.301517

Bowen, B., Steinberg, J., Laemmli, U. K., & Weintraub, H. (1980). The detection of DNA-binding proteins by protein blotting. Nucleic Acids Research, 8(1), 1–20.

Bozdech, Z., Llinás, M., Pulliam, B. L., Wong, E. D., Zhu, J., & DeRisi, J. L. (2003a). The Transcriptome of the Intraerythrocytic Developmental Cycle of Plasmodium falciparum. PLOS Biology, 1(1), e5. 10.1371/journal.pbio.0000005

Bozdech, Z., Llinás, M., Pulliam, B. L., Wong, E. D., Zhu, J., & DeRisi, J. L. (2003b). The Transcriptome of the Intraerythrocytic Developmental Cycle of Plasmodium falciparum. PLoS Biology, 1(1), e5. 10.1371/journal.pbio.0000005

Breiling, A., & Lyko, F. (2015). Epigenetic regulatory functions of DNA modifications: 5-methylcytosine and beyond. Epigenetics & Chromatin, 8(1), 24. 10.1186/s13072-015-0016-6

Bunnik, E. M., Chung, D.-W. D., Hamilton, M., Ponts, N., Saraf, A., Prudhomme, J., Florens, L., & Le Roch, K. G. (2013). Polysome profiling reveals translational control of gene expression in the human malaria parasite Plasmodium falciparum. Genome Biology, 14(11), R128. 10.1186/gb-2013-14-11-r128

Cadet, J., Douki, T., & Ravanat, J.-L. (2010). Oxidatively generated base damage to cellular DNA. Free Radical Biology and Medicine, 49(1), 9–21. 10.1016/j.freeradbiomed.2010.03.025

Coetzee, N., Sidoli, S., van Biljon, R., Painter, H., Llinás, M., Garcia, B. A., & Birkholtz, L.-M. (2017). Quantitative chromatin proteomics reveals a dynamic histone post-translational modification landscape that defines asexual and sexual Plasmodium falciparum parasites. Scientific Reports, 7(1), 607. 10.1038/s41598-017-00687-7

Comeaux, C. A., & Duraisingh, M. T. (2007). Unravelling a histone code for malaria virulence. Molecular Microbiology, 66(6), 1291–1295. 10.1111/j.1365-2958.2007.06038.x

Connacher, J., Josling, G. A., Orchard, L. M., Reader, J., Llinás, M., & Birkholtz, L.-M. (2021). H3K36 methylation reprograms gene expression to drive early gametocyte development in Plasmodium falciparum. Epigenetics & Chromatin, 14(1), 19. 10.1186/s13072-021-00393-9

Crowley, V. M., Rovira-Graells, N., de Pouplana, L. R., & Cortés, A. (2011). Heterochromatin formation in bistable chromatin domains controls the epigenetic repression of clonally variant Plasmodium falciparum genes linked to erythrocyte invasion. Molecular Microbiology, 80(2), 391–406. 10.1111/j.1365-2958.2011.07574.x

Cui, L., Lindner, S., & Miao, J. (2015a). Translational regulation during stage transitions in malaria parasites. Annals of the New York Academy of Sciences, 1342(1), 1–9. 10.1111/nyas.12573

Cui, L., Lindner, S., & Miao, J. (2015b). Translational regulation during stage transitions in malaria parasites. Annals of the New York Academy of Sciences, 1342(1), 1–9. 10.1111/nyas.12573

Cui, L., & Miao, J. (2010). Chromatin-Mediated Epigenetic Regulation in the Malaria Parasite Plasmodium falciparum. Eukaryotic Cell, 9(8), 1138–1149. 10.1128/EC.00036-10

David, S. S., O’Shea, V. L., & Kundu, S. (2007). Base-excision repair of oxidative DNA damage. Nature, 447(7147), 941–950. 10.1038/nature05978

Dizdaroglu, M. (2002, May 1). Formation of 8-hydroxyguanine moiety in deoxyribonucleic acid on .gamma*.-*irradiation in aqueous solution [Research-article]. American Chemical Society. (world). ACS Publications. 10.1021/bi00337a032

Dobin, A., Davis, C. A., Schlesinger, F., Drenkow, J., Zaleski, C., Jha, S., Batut, P., Chaisson, M., & Gingeras, T. R. (2013). STAR: Ultrafast universal RNA-seq aligner. Bioinformatics, 29(1), 15–21. 10.1093/bioinformatics/bts635

Doerig, C., Rayner, J. C., Scherf, A., & Tobin, A. B. (2015). Post-translational protein modifications in malaria parasites. Nature Reviews. Microbiology, 13(3), 160–172. 10.1038/nrmicro3402

Duffy, M. F., Selvarajah, S. A., Josling, G. A., & Petter, M. (2014). Epigenetic regulation of the Plasmodium falciparum genome. Briefings in Functional Genomics, 13(3), 203–216. 10.1093/bfgp/elt047

Fedeles, B. I. (2017). G-quadruplex-forming promoter sequences enable transcriptional activation in response to oxidative stress. Proceedings of the National Academy of Sciences of the United States of America, 114(11), 2788–2790. 10.1073/pnas.1701244114

Fleming, A. M., & Burrows, C. J. (2020). Interplay of Guanine Oxidation and G-Quadruplex Folding in Gene Promoters. Journal of the American Chemical Society, 142(3), 1115–1136. 10.1021/jacs.9b11050

Fleming, A. M., & Burrows, C. J. (2021). Oxidative stress-mediated epigenetic regulation by G-quadruplexes. NAR Cancer, 3(3), zcab038. 10.1093/narcan/zcab038

Fleming, A. M., Ding, Y., & Burrows, C. J. (2017). Oxidative DNA damage is epigenetic by regulating gene transcription via base excision repair. Proceedings of the National Academy of Sciences, 114(10), 2604–2609. 10.1073/pnas.1619809114

Fleming, A. M., Zhu, J., Ding, Y., & Burrows, C. J. (2017). 8-Oxo-7,8-dihydroguanine in the Context of a Gene Promoter G-Quadruplex Is an On–Off Switch for Transcription. ACS Chemical Biology, 12(9), 2417–2426. 10.1021/acschembio.7b00636

Francis, S. E., David J. Sullivan Jr, & Goldberg, and D. E. (1997). HEMOGLOBIN METABOLISM IN THE MALARIA PARASITE PLASMODIUM FALCIPARUM. Annual Review of Microbiology, 51(Volume 51, 1997), 97–123. 10.1146/annurev.micro.51.1.97

Fukuda, K., Yoshida, H., Sato, T., Furumoto, T., Mizutani-Koseki, Y., Suzuki, Y., Saito, Y., Takemori, T., Kimura, M., Sato, H., Taniguchi, M., Nishikawa, S., Nakayama, T., & Koseki, H. (2003). Mesenchymal expression of Foxl1, a winged helix transcriptional factor, regulates generation and maintenance of gut-associated lymphoid organs. Developmental Biology, 255(2), 278–289. 10.1016/s0012-1606(02)00088-x

Gardner, M. J., Hall, N., Fung, E., White, O., Berriman, M., Hyman, R. W., Carlton, J. M., Pain, A., Nelson, K. E., Bowman, S., Paulsen, I. T., James, K., Eisen, J. A., Rutherford, K., Salzberg, S. L., Craig, A., Kyes, S., Chan, M.-S., Nene, V., … Barrell, B. (2002a). Genome sequence of the human malaria parasite Plasmodium falciparum. Nature, 419(6906),. 10.1038/nature01097

Gardner, M. J., Hall, N., Fung, E., White, O., Berriman, M., Hyman, R. W., Carlton, J. M., Pain, A., Nelson, K. E., Bowman, S., Paulsen, I. T., James, K., Eisen, J. A., Rutherford, K., Salzberg, S. L., Craig, A., Kyes, S., Chan, M.-S., Nene, V., … Barrell, B. (2002b). Genome sequence of the human malaria parasite Plasmodium falciparum. Nature, 419(6906), 10.1038/nature01097.

Gazanion, E., Lacroix, L., Alberti, P., Gurung, P., Wein, S., Cheng, M., Mergny, J.-L., Gomes, A. R., & Lopez-Rubio, J.-J. (2020). Genome wide distribution of G-quadruplexes and their impact on gene expression in malaria parasites. PLoS Genetics, 16(7), e1008917. 10.1371/journal.pgen.1008917

Gillespie, M. N., & Wilson, G. L. (2007). Bending and breaking the code: Dynamic changes in promoter integrity may underlie a new mechanism regulating gene expression. American Journal of Physiology-Lung Cellular and Molecular Physiology, 292(1), L1–L3. 10.1152/ajplung.00275.2006

Gissot, M., Choi, S.-W., Thompson, R. F., Greally, J. M., & Kim, K. (2008). Toxoplasma gondii and Cryptosporidium parvum Lack Detectable DNA Cytosine Methylation. Eukaryotic Cell, 7(3), 537–540. 10.1128/EC.00448-07

Goffeau, A., Barrell, B. G., Bussey, H., Davis, R. W., Dujon, B., Feldmann, H., Galibert, F., Hoheisel, J. D., Jacq, C., Johnston, M., Louis, E. J., Mewes, H. W., Murakami, Y., Philippsen, P., Tettelin, H., & Oliver, S. G. (1996). Life with 6000 genes. Science, 274(5287), 546, 563–567. 10.1126/science.274.5287.546

Gómez-Díaz, E., Yerbanga, R. S., Lefèvre, T., Cohuet, A., Rowley, M. J., Ouedraogo, J. B., & Corces, V. G. (2017). Epigenetic regulation of Plasmodium falciparum clonally variant gene expression during development in Anopheles gambiae. Scientific Reports, 7(1), 40655. 10.1038/srep40655

Gorini, F., Ambrosio, S., Lania, L., Majello, B., & Amente, S. (2023). The Intertwined Role of 8-oxodG and G4 in Transcription Regulation. International Journal of Molecular Sciences, 24(3), 2031. 10.3390/ijms24032031

Gorini, F., Scala, G., Ambrosio, S., Majello, B., & Amente, S. (2022). OxiDIP-Seq for Genome-wide Mapping of Damaged DNA Containing 8-Oxo-2’-Deoxyguanosine. Bio-Protocol, 12(21), e4540. 10.21769/BioProtoc.4540

Gorini, F., Scala, G., Di Palo, G., Dellino, G. I., Cocozza, S., Pelicci, P. G., Lania, L., Majello, B., & Amente, S. (2020). The genomic landscape of 8-oxodG reveals enrichment at specific inherently fragile promoters. Nucleic Acids Research, 48(8), 4309–4324. 10.1093/nar/gkaa175

Gupta, A. P., & Bozdech, Z. (2017). Epigenetic landscapes underlining global patterns of gene expression in the human malaria parasite, *Plasmodium falciparum. International Journal for Parasitology*, Singapore Malaria Network Meeting (SingMalNet) 2016, 47(7), 399–407. 10.1016/j.ijpara.2016.10.008

Gupta, A. P., Zhu, L., Tripathi, J., Kucharski, M., Patra, A., & Bozdech, Z. (2017). Histone 4 lysine 8 acetylation regulates proliferation and host–pathogen interaction in Plasmodium falciparum. Epigenetics & Chromatin, 10(1), 40. 10.1186/s13072-017-0147-z

Hack, L. M., Dick, A. L. W., & Provençal, N. (2016). Epigenetic mechanisms involved in the effects of stress exposure: Focus on 5-hydroxymethylcytosine: Table 1: Environmental Epigenetics, 2(3), dvw016. 10.1093/eep/dvw016

Hahm, J. Y., Park, J., Jang, E.-S., & Chi, S. W. (2022). 8-Oxoguanine: From oxidative damage to epigenetic and epitranscriptional modification. Experimental & Molecular Medicine, 54(10), 1626–1642. 10.1038/s12276-022-00822-z

Haltiwanger, B. M., Matsumoto, Y., Nicolas, E., Dianov, G. L., Bohr, V. A., & Taraschi, T. F. (2000). DNA base excision repair in human malaria parasites is predominantly by a long-patch pathway. Biochemistry, 39(4), 763–772. 10.1021/bi9923151

Hammam, E., Ananda, G., Sinha, A., Scheidig-Benatar, C., Bohec, M., Preiser, P. R., Dedon, P. C., Scherf, A., & Vembar, S. S. (2020). Discovery of a new predominant cytosine DNA modification that is linked to gene expression in malaria parasites. Nucleic Acids Research, 48(1), 184–199. 10.1093/nar/gkz1093

Hazra, T. K., Das, A., Das, S., Choudhury, S., Kow, Y. W., & Roy, R. (2007). Oxidative DNA damage repair in mammalian cells: A new perspective. *DNA Repair*, Repair of Small Base Lesions in DNA—from Molecular Biology to Phenotype, 6(4), 470–480. 10.1016/j.dnarep.2006.10.011

Hegde, M. L., Hazra, T. K., & Mitra, S. (2008). Early steps in the DNA base excision/single-strand interruption repair pathway in mammalian cells. Cell Research, 18(1), 27–47. 10.1038/cr.2008.8

Herrera-Solorio, A. M., Vembar, S. S., MacPherson, C. R., Lozano-Amado, D., Meza, G. R., Xoconostle-Cazares, B., Martins, R. M., Chen, P., Vargas, M., Scherf, A., & Hernández-Rivas, R. (2019). Clipped histone H3 is integrated into nucleosomes of DNA replication genes in the human malaria parasite Plasmodium falciparum. EMBO Reports, 20(4), e46331. 10.15252/embr.201846331

Hughes, K. R., Philip, N., Starnes, G. L., Taylor, S., & Waters, A. P. (2010). From cradle to grave: RNA biology in malaria parasites. Wiley Interdisciplinary Reviews. RNA, 1(2), 287–303. 10.1002/wrna.30

Issar, N., Ralph, S. A., Mancio-Silva, L., Keeling, C., & Scherf, A. (2009). Differential sub-nuclear localisation of repressive and activating histone methyl modifications in *P. falciparum*. Microbes and Infection, 11(3), 403–407. 10.1016/j.micinf.2008.12.010

Jiang, L., Mu, J., Zhang, Q., Ni, T., Srinivasan, P., Rayavara, K., Yang, W., Turner, L., Lavstsen, T., Theander, T. G., Peng, W., Wei, G., Jing, Q., Wakabayashi, Y., Bansal, A., Luo, Y., Ribeiro, J. M. C., Scherf, A., Aravind, L., … Miller, L. H. (2013). PfSETvs methylation of histone H3K36 represses virulence genes in Plasmodium falciparum. Nature, 499(7457), 223–227. 10.1038/nature12361

Karmodiya, K., Pradhan, S. J., Joshi, B., Jangid, R., Reddy, P. C., & Galande, S. (2015). A comprehensive epigenome map of Plasmodium falciparum reveals unique mechanisms of transcriptional regulation and identifies H3K36me2 as a global mark of gene suppression. Epigenetics & Chromatin, 8, 32. 10.1186/s13072-015-0029-1

Kaur, I., Zeeshan, M., Saini, E., Kaushik, A., Mohmmed, A., Gupta, D., & Malhotra, P. (2016). Widespread occurrence of lysine methylation in Plasmodium falciparum proteins at asexual blood stages. Scientific Reports, 6(1), 35432. 10.1038/srep35432

Kensche, P. R., Hoeijmakers, W. A. M., Toenhake, C. G., Bras, M., Chappell, L., Berriman, M., & Bártfai, R. (2016). The nucleosome landscape of Plasmodium falciparum reveals chromatin architecture and dynamics of regulatory sequences. Nucleic Acids Research, 44(5), 2110–2124. 10.1093/nar/gkv1214

Kino, K., Hirao-Suzuki, M., Morikawa, M., Sakaga, A., & Miyazawa, H. (2017). Generation, repair and replication of guanine oxidation products. Genes and Environment, 39, 21. 10.1186/s41021-017-0081-0

Klose, R. J., & Bird, A. P. (2006). Genomic DNA methylation: The mark and its mediators. Trends in Biochemical Sciences, 31(2), 89–97. 10.1016/j.tibs.2005.12.008

Krueger, F., James, F., Ewels, P., Afyounian, E., Weinstein, M., Schuster-Boeckler, B., Hulselmans, G., & sclamons. (2023). FelixKrueger/TrimGalore: V0.6.10 - add default decompression path (Version 0.6.10) [Computer software]. Zenodo. 10.5281/zenodo.7598955

Kumar, M., Skillman, K., & Duraisingh, M. T. (2021). Linking nutrient sensing and gene expression in Plasmodium falciparum blood-stage parasites. Molecular Microbiology, 115(5), 891–900. 10.1111/mmi.14652

Kumar, S., Chinnusamy, V., & Mohapatra, T. (2018). Epigenetics of Modified DNA Bases: 5-Methylcytosine and Beyond. Frontiers in Genetics, 9, 640. 10.3389/fgene.2018.00640

Kuznetsova, A. A., Fedorova, O. S., & Kuznetsov, N. A. (2020). Lesion Recognition and Cleavage of Damage-Containing Quadruplexes and Bulged Structures by DNA Glycosylases. Frontiers in Cell and Developmental Biology, 8, 595687. 10.3389/fcell.2020.595687

Le Roch, K. G., Johnson, J. R., Florens, L., Zhou, Y., Santrosyan, A., Grainger, M., Yan, S. F., Williamson, K. C., Holder, A. A., Carucci, D. J., Yates, J. R., & Winzeler, E. A. (2004). Global analysis of transcript and protein levels across the Plasmodium falciparum life cycle. Genome Research, 14(11), 2308–2318. 10.1101/gr.2523904

Li, H., & Durbin, R. (2010). Fast and accurate long-read alignment with Burrows–Wheeler transform. Bioinformatics, 26(5), 589–595. 10.1093/bioinformatics/btp698

Li, H., Handsaker, B., Wysoker, A., Fennell, T., Ruan, J., Homer, N., Marth, G., Abecasis, G., & Durbin, R. (2009). The Sequence Alignment/Map format and SAMtools. Bioinformatics, 25(16), 2078–2079. 10.1093/bioinformatics/btp352

López-Barragán, M. J., Lemieux, J., Quiñones, M., Williamson, K. C., Molina-Cruz, A., Cui, K., Barillas-Mury, C., Zhao, K., & Su, X. (2011). Directional gene expression and antisense transcripts in sexual and asexual stages of Plasmodium falciparum. BMC Genomics, 12(1), 587. 10.1186/1471-2164-12-587

Lu, T., Pan, Y., Kao, S.-Y., Li, C., Kohane, I., Chan, J., & Yankner, B. A. (2004). Gene regulation and DNA damage in the ageing human brain. Nature, 429(6994), 883–891. 10.1038/nature02661

Lucky, A. B., Wang, C., Li, X., Chim-Ong, A., Adapa, S. R., Quinlivan, E. P., Jiang, R., Cui, L., & Miao, J. (2023). Characterization of the dual role of Plasmodium falciparum DNA methyltransferase in regulating transcription and translation. Nucleic Acids Research, 51(8), 3918–3933. 10.1093/nar/gkad248

Mayr, C. (2017). Regulation by 3′-Untranslated Regions. Annual Review of Genetics, 51(Volume 51, 2017), 171–194. 10.1146/annurev-genet-120116-024704

Neeley, W. L., & Essigmann, J. M. (2006). Mechanisms of Formation, Genotoxicity, and Mutation of Guanine Oxidation Products. Chemical Research in Toxicology, 19(4), 491–505. 10.1021/tx0600043

Ngwa, C. J., Gross, M. R., Musabyimana, J.-P., Pradel, G., & Deitsch, K. W. (2021). The Role of the Histone Methyltransferase PfSET10 in Antigenic Variation by Malaria Parasites: A Cautionary Tale. mSphere, 6(1), 10.1128/msphere.01217-20.

Ngwa, C. J., Kiesow, M. J., Papst, O., Orchard, L. M., Filarsky, M., Rosinski, A. N., Voss, T. S., Llinás, M., & Pradel, G. (2017). Transcriptional Profiling Defines Histone Acetylation as a Regulator of Gene Expression during Human-to-Mosquito Transmission of the Malaria Parasite Plasmodium falciparum. Frontiers in Cellular and Infection Microbiology, 7. 10.3389/fcimb.2017.00320

Nowling, R. J., Beal, C. R., Emrich, S., Behura, S. K., Halfon, M. S., & Duman-Scheel, M. (2020). PeakMatcher: Matching Peaks Across Genome Assemblies. Proceedings of the 11th ACM International Conference on Bioinformatics, Computational Biology and Health Informatics, BCB’20, 1. 10.1145/3388440.3414907

Obermann, T., Sakshaug, T., Kanagaraj, V. V., Abentung, A., de Sousa, M. M. L., Hagen, L., Sarno, A., Bjørås, M., & Scheffler, K. (2025). Genomic 8-oxoguanine modulates gene transcription independent of its repair by DNA glycosylases OGG1 and MUTYH. Redox Biology, 79, 103461. 10.1016/j.redox.2024.103461

Painter, H. J., Chung, N. C., Sebastian, A., Albert, I., Storey, J. D., & Llinás, M. (2018). Genome-wide real-time in vivo transcriptional dynamics during Plasmodium falciparum blood-stage development. Nature Communications, 9(1), 2656. 10.1038/s41467-018-04966-3

Pan, L., Hao, W., Xue, Y., Wang, K., Zheng, X., Luo, J., Ba, X., Xiang, Y., Qin, X., Bergwik, J., Tanner, L., Egesten, A., Brasier, A. R., & Boldogh, I. (2023a). 8-Oxoguanine targeted by 8-oxoguanine DNA glycosylase 1 (OGG1) is central to fibrogenic gene activation upon lung injury. Nucleic Acids Research, 51(3), 1087–1102. 10.1093/nar/gkac1241

Pan, L., Hao, W., Xue, Y., Wang, K., Zheng, X., Luo, J., Ba, X., Xiang, Y., Qin, X., Bergwik, J., Tanner, L., Egesten, A., Brasier, A. R., & Boldogh, I. (2023b). 8-Oxoguanine targeted by 8-oxoguanine DNA glycosylase 1 (OGG1) is central to fibrogenic gene activation upon lung injury. Nucleic Acids Research, 51(3), 1087–1102. 10.1093/nar/gkac1241

Pan, L., Hao, W., Zheng, X., Zeng, X., Ahmed Abbasi, A., Boldogh, I., & Ba, X. (2017). OGG1-DNA interactions facilitate NF-κB binding to DNA targets. Scientific Reports, 7(1), 43297. 10.1038/srep43297

Pan, L., Vlahopoulos, S., Tanner, L., Bergwik, J., Bacsi, A., Radak, Z., Egesten, A., Ba, X., Brasier, A. R., & Boldogh, I. (2023). Substrate-specific binding of 8-oxoguanine DNA glycosylase 1 (OGG1) reprograms mucosal adaptations to chronic airway injury. Frontiers in Immunology, 14. 10.3389/fimmu.2023.1186369

Pan, L., Zhu, B., Hao, W., Zeng, X., Vlahopoulos, S. A., Hazra, T. K., Hegde, M. L., Radak, Z., Bacsi, A., Brasier, A. R., Ba, X., & Boldogh, I. (2016a). Oxidized Guanine Base Lesions Function in 8-Oxoguanine DNA Glycosylase-1-mediated Epigenetic Regulation of Nuclear Factor κB-driven Gene Expression *. Journal of Biological Chemistry, 291(49), 25553–25566. 10.1074/jbc.M116.751453

Pan, L., Zhu, B., Hao, W., Zeng, X., Vlahopoulos, S. A., Hazra, T. K., Hegde, M. L., Radak, Z., Bacsi, A., Brasier, A. R., Ba, X., & Boldogh, I. (2016b). Oxidized Guanine Base Lesions Function in 8-Oxoguanine DNA Glycosylase-1-mediated Epigenetic Regulation of Nuclear Factor κB-driven Gene Expression *. Journal of Biological Chemistry, 291(49), 25553–25566. 10.1074/jbc.M116.751453

Pastukh, V., Roberts, J. T., Clark, D. W., Bardwell, G. C., Patel, M., Al-Mehdi, A.-B., Borchert, G. M., & Gillespie, M. N. (2015). An oxidative DNA “damage” and repair mechanism localized in the VEGF promoter is important for hypoxia-induced VEGF mRNA expression. American Journal of Physiology-Lung Cellular and Molecular Physiology, 309(11), L1367–L1375. 10.1152/ajplung.00236.2015

Ponts, N., Fu, L., Harris, E. Y., Zhang, J., Chung, D.-W. D., Cervantes, M. C., Prudhomme, J., Atanasova-Penichon, V., Zehraoui, E., Bunnik, E. M., Rodrigues, E. M., Lonardi, S., Hicks, G. R., Wang, Y., & Le Roch, K. G. (2013). Genome-wide mapping of DNA methylation in the human malaria parasite Plasmodium falciparum. Cell Host & Microbe, 14(6), 696–706. 10.1016/j.chom.2013.11.007

Quinlan, A. R., & Hall, I. M. (2010). BEDTools: A flexible suite of utilities for comparing genomic features. Bioinformatics, 26(6), 841–842. 10.1093/bioinformatics/btq033

Ramírez, F., Dündar, F., Diehl, S., Grüning, B. A., & Manke, T. (2014). deepTools: A flexible platform for exploring deep-sequencing data. Nucleic Acids Research, 42(Web Server issue), W187-191. 10.1093/nar/gku365

Sampath, H., & Lloyd, R. S. (2019). Roles of OGG1 in Transcriptional Regulation and Maintenance of Metabolic Homeostasis. DNA Repair, 81, 102667. 10.1016/j.dnarep.2019.102667

Saraf, A., Cervantes, S., Bunnik, E. M., Ponts, N., Sardiu, M. E., Chung, D.-W. D., Prudhomme, J., Varberg, J. M., Wen, Z., Washburn, M. P., Florens, L., & Le Roch, K. G. (2016). Dynamic and Combinatorial Landscape of Histone Modifications during the Intraerythrocytic Developmental Cycle of the Malaria Parasite. Journal of Proteome Research, 15(8), 2787–2801. 10.1021/acs.jproteome.6b00366

Shikata, M., Matsuda, Y., Ando, K., Nishii, A., Takemura, M., Yokota, A., & Kohchi, T. (2004). Characterization of Arabidopsis ZIM, a member of a novel plant-specific GATA factor gene family. Journal of Experimental Botany, 55(397), 631–639. 10.1093/jxb/erh078

Sun, W., Guan, M., & Li, X. (2014). 5-Hydroxymethylcytosine-Mediated DNA Demethylation in Stem Cells and Development. Stem Cells and Development, 23(9), 923–930. 10.1089/scd.2013.0428

Tian, B., & Manley, J. L. (2017). Alternative polyadenylation of mRNA precursors. Nature Reviews Molecular Cell Biology, 18(1), 18–30. 10.1038/nrm.2016.116

Tiwari, A., Kuldeep, J., Siddiqi, M. I., & Habib, S. (2020). Plasmodium falciparum Apn1 homolog is a mitochondrial base excision repair protein with restricted enzymatic functions. The FEBS Journal, 287(3), 589–606. 10.1111/febs.15032

Tiwari, A., Verma, N., Shukla, H., Mishra, S., Kennedy, K., Chatterjee, T., Kuldeep, J., Parwez, S., Siddiqi, M. I., Ralph, S. A., Mishra, S., & Habib, S. (2024). DNA N-glycosylases Ogg1 and EndoIII as components of base excision repair in Plasmodium falciparum organelles.International Journal for Parasitology, 54(13), 675–689. 10.1016/j.ijpara.2024.06.005

Toenhake, C. G., Fraschka, S. A.-K., Vijayabaskar, M. S., Westhead, D. R., van Heeringen, S. J., & Bártfai, R. (2018). Chromatin Accessibility-Based Characterization of the Gene Regulatory Network Underlying Plasmodium falciparum Blood-Stage Development. Cell Host & Microbe, 23(4), 557–569.e9. 10.1016/j.chom.2018.03.007

Trager, W., & Jensen, J. B. (1976). Human malaria parasites in continuous culture. *Science (New York*, N.Y.), 193(4254), 673–675. 10.1126/science.781840

Trelle, M. B., Salcedo-Amaya, A. M., Cohen, A. M., Stunnenberg, H. G., & Jensen, O. N. (2009). Global Histone Analysis by Mass Spectrometry Reveals a High Content of Acetylated Lysine Residues in the Malaria Parasite Plasmodium falciparum. Journal of Proteome Research, 8(7), 3439–3450. 10.1021/pr9000898

van Biljon, R., van Wyk, R., Painter, H. J., Orchard, L., Reader, J., Niemand, J., Llinás, M., & Birkholtz, L.-M. (2019). Hierarchical transcriptional control regulates Plasmodium falciparum sexual differentiation. BMC Genomics, 20(1), 920. 10.1186/s12864-019-6322-9

Vembar, S. S., Droll, D., & Scherf, A. (2016). Translational regulation in blood stages of the malaria parasite Plasmodium spp.: Systems-wide studies pave the way. WIREs RNA, 7(6), 772–792. 10.1002/wrna.1365

Verma, N., Shukla, H., Tiwari, A., Mishra, S., & Habib, S. (2021). *Plasmodium* Ape1 is a multifunctional enzyme in mitochondrial base excision repair and is required for efficient transition from liver to blood stage infection. DNA Repair, 101, 103098. 10.1016/j.dnarep.2021.103098

Visnes, T., Cázares-Körner, A., Hao, W., Wallner, O., Masuyer, G., Loseva, O., Mortusewicz, O., Wiita, E., Sarno, A., Manoilov, A., Astorga-Wells, J., Jemth, A.-S., Pan, L., Sanjiv, K., Karsten, S., Gokturk, C., Grube, M., Homan, E. J., Hanna, B. M. F., … Helleday, T. (2018). Small-molecule inhibitor of OGG1 suppresses proinflammatory gene expression and inflammation. Science, 362(6416), 834–839. 10.1126/science.aar8048

Voss, T. S., Bozdech, Z., & Bártfai, R. (2014). Epigenetic memory takes center stage in the survival strategy of malaria parasites. Current Opinion in Microbiology, 20, 88–95. 10.1016/j.mib.2014.05.007

Wickham, H. (2016) ggplot2 Elegant Graphics for Data Analysis. Springer-Verlag, New York. - References—Scientific Research Publishing. (n.d.). Retrieved June 18, 2026, from https://www.scirp.org/reference/referencespapers?referenceid=3610100

Wilcoxon Signed Rank Test for Paired Comparisons of Clustered Data | Biometrics | Oxford Academic. (n.d.). Retrieved June 4, 2026, from https://academic.oup.com/biometrics/article-abstract/62/1/185/7306230?redirectedFrom=fulltext

Witmer, K., Fraschka, S. A., Vlachou, D., Bártfai, R., & Christophides, G. K. (2020). An epigenetic map of malaria parasite development from host to vector. Scientific Reports, 10(1), 6354. 10.1038/s41598-020-63121-5

Young, J. A., Fivelman, Q. L., Blair, P. L., de la Vega, P., Le Roch, K. G., Zhou, Y., Carucci, D. J., Baker, D. A., & Winzeler, E. A. (2005). The *Plasmodium falciparum* sexual development transcriptome: A microarray analysis using ontology-based pattern identification. Molecular and Biochemical Parasitology, 143(1), 67–79. 10.1016/j.molbiopara.2005.05.007

Zhang, M., Joyce, B. R., Sullivan, W. J., & Nussenzweig, V. (2013). Translational control in Plasmodium and toxoplasma parasites. Eukaryotic Cell, 12(2), 161–167. 10.1128/EC.00296-12

Zhang, Y., Liu, T., Meyer, C. A., Eeckhoute, J., Johnson, D. S., Bernstein, B. E., Nusbaum, C., Myers, R. M., Brown, M., Li, W., & Liu, X. S. (2008). Model-based Analysis of ChIP-Seq (MACS). Genome Biology, 9(9), R137. 10.1186/gb-2008-9-9-r137

Zhao, Y., Wang, J., Liang, F., Liu, Y., Wang, Q., Zhang, H., Jiang, M., Zhang, Z., Zhao, W., Bao, Y., Zhang, Z., Wu, J., Asmann, Y. W., Li, R., & Xiao, J. (2019). NucMap: A database of genome-wide nucleosome positioning map across species. Nucleic Acids Research, 47(D1), D163–D169. 10.1093/nar/gky980

Zhong, Y., Zhang, X., Feng, R., Fan, Y., Zhang, Z., Zhang, Q.-W., Wan, J.-B., Wang, Y., Yu, H., & Li, G. (2024). OGG1: An emerging multifunctional therapeutic target for the treatment of diseases caused by oxidative DNA damage. Medicinal Research Reviews, 44(6), 2825–2848. 10.1002/med.22068

Zhou, J., Liu, M., Fleming, A. M., Burrows, C. J., & Wallace, S. S. (2013). Neil3 and NEIL1 DNA Glycosylases Remove Oxidative Damages from Quadruplex DNA and Exhibit Preferences for Lesions in the Telomeric Sequence Context. The Journal of Biological Chemistry, 288(38), 27263–27272. 10.1074/jbc.M113.479055

