## Supplementary Info for "Genomic 8-oxoguanine is associated with transcriptionally active chromatin and elevated gene expression in *Plasmodium falciparum*": DiA_20260616_8_oxoG_Supplementary_information.docx

**Supplementary information for Acharya and Vembar, 2026**

**This document contains:**

Supplementary Figures S1 to S7, with legends

**The following are provided as Excel sheets:**

**Supplementary Table S1:** Mapping statistics of the fastq files generated from OxiDIP-seq and RNA-seq

**Supplementary Table S2:** MACS2-derived 8-oxoG-containing peaks identified for each timepoint by OxiDiP-seq

**Supplementary Table S3: Genes marked with 8-oxoG at different timepoints**

**Supplementary Table S4:** HTSeq-generated raw read counts and normalised RPKM data for all *P. falciparum* genes in the in-house 10 hpi transcriptomic datasets used in this study

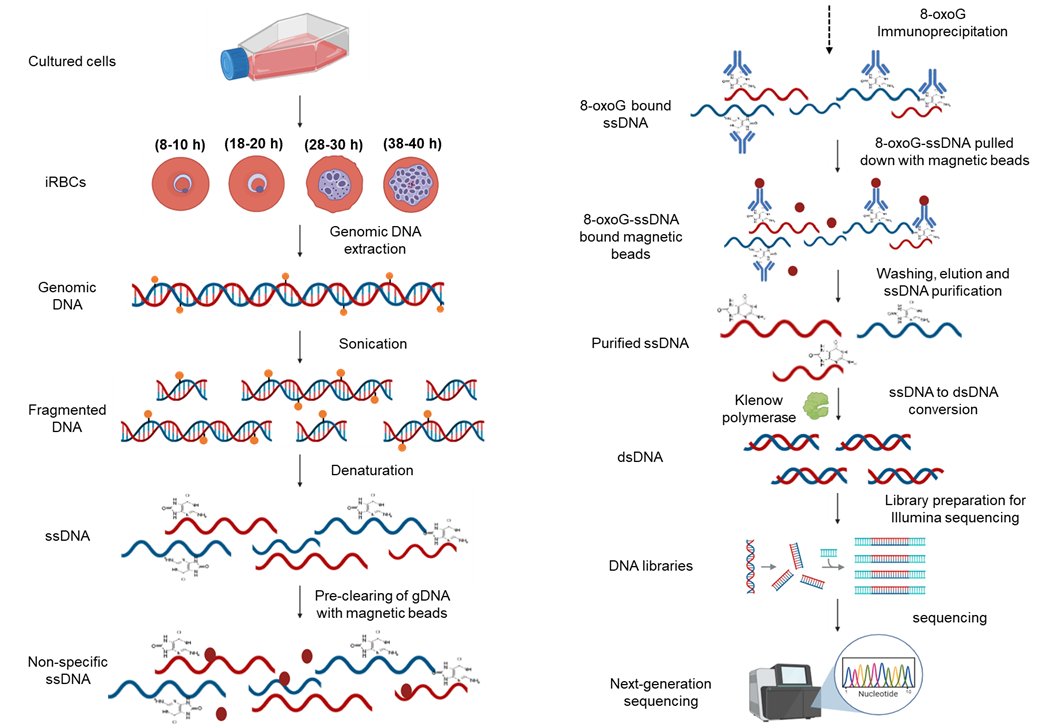

**Supplementary Figure S1**: Detailed overview of the OxiDIP-seq workflow (Adapted from Gorini et al., 2022). Image generated using BioRender.

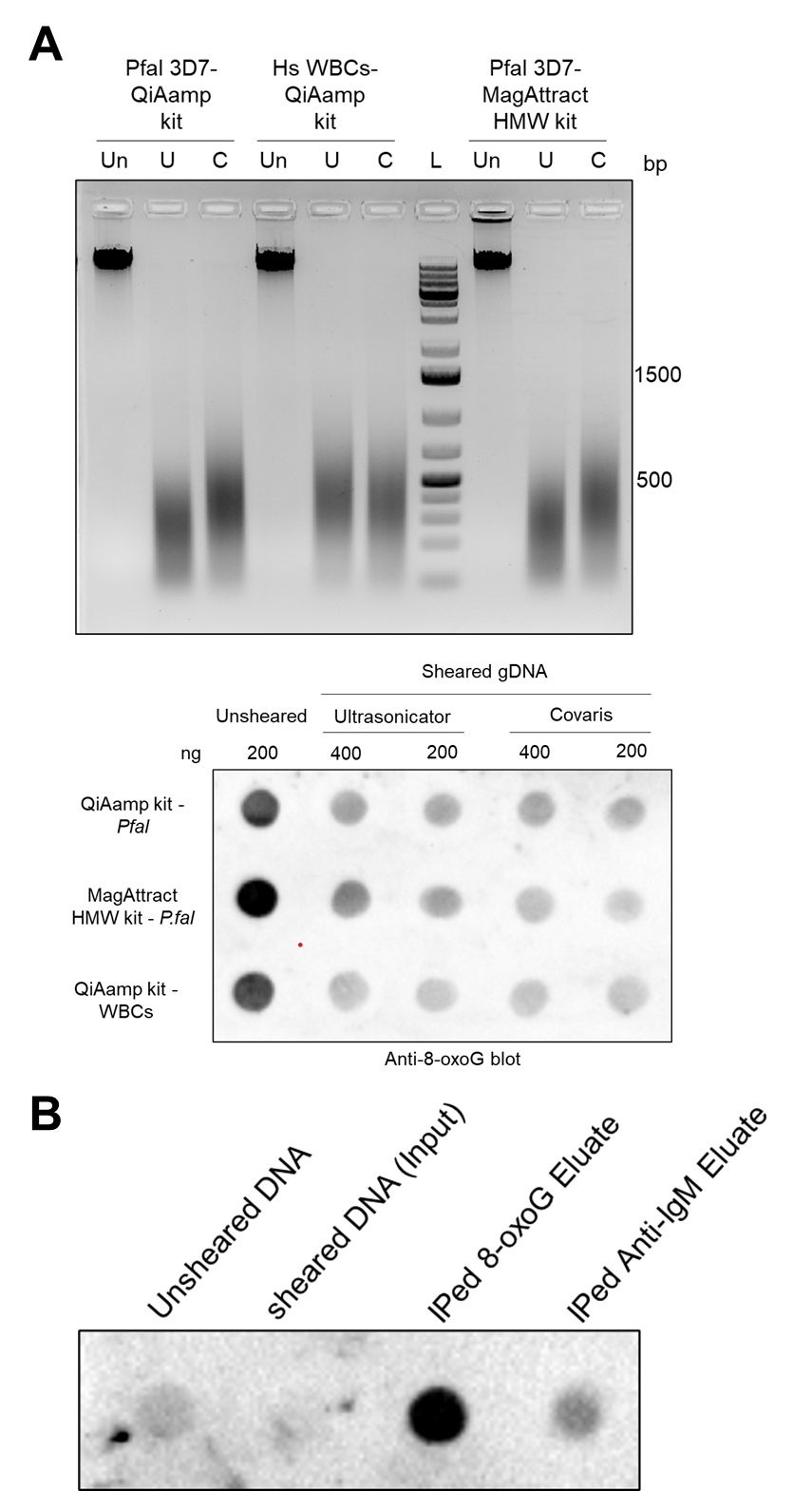

**Supplementary Figure S2:** **Optimization of shearing for *P. falciparum* genomic DNA followed by 8-oxoG detection.** **(A)** ***Upper panel:*** Comparative agarose gel electrophoresis of genomic DNA (gDNA) prepared using the indicated kit and sheared using the Covaris S2 ultrasonicator (C) or bath ultrasonicator (U). Pfal = *P. falciparum*; Hs = human. L = 1 kb plus DNA ladder (Thermo Fisher Scientific). ***Lower panel:*** Anti-8-oxoG dot blot analysis of the sheared gDNA. Un = Unsheared DNA. **(B)** Dot blot of 8-oxoG-containing gDNA enriched using the optimised OxiDIP protocol. After immunoprecipitation, 25 ng of eluate or input gDNA (unsheared or sheared) was subjected to anti-8-oxoG South-western blotting.

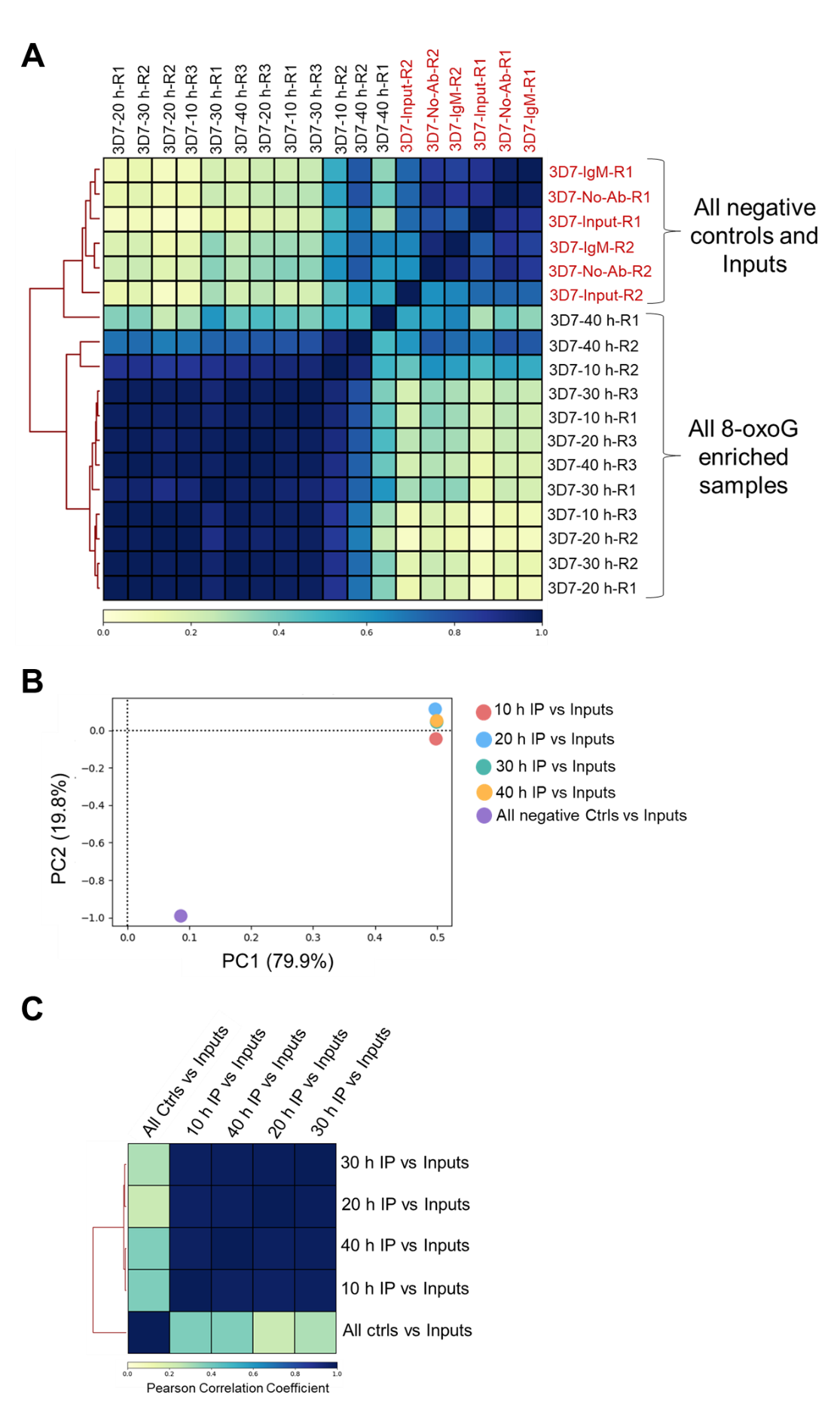

**Supplementary Figure S3:** **Quality control and correlation analysis of 8-oxoG enrichment profiles relative to controls. (A)** Heatmap displaying pairwise Pearson Correlation Coefficient (PCC) R between the indicated OxiDIP-seq samples. Correlations were computed using BAM alignment files derived from sequencing data mapped to the *P. falciparum* 3D7 reference genome. The colour gradient represents the strength of correlation, with values ranging from 0 (no correlation) to 1 (perfect correlation). **(B)** Principal Component Analysis of the different MACS2-derived fold enrichment (FE) profiles of OxiDIP-seq samples relative to Input DNA was performed using the plotPCA function of DeepTools on a multibigwigsummary file. **(C)** Heatmap displaying pairwise PCC between the indicated OxiDIP-seq profiles. Correlations were computed using FE.bedgraph files derived from MACS2 analysis. The color scale indicates values of R from 0 to 1.

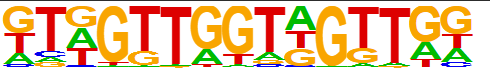

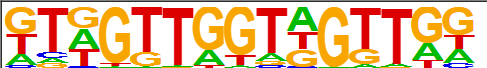

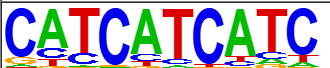

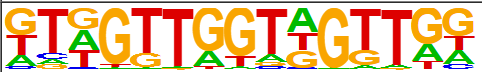

10 h

20 h

30 h

40 h

**Supplementary Figure S4: Stage-specific motif enrichment within 8-oxoG-containing peaks.** Motif enrichment analysis of 8-oxoG-containing peaks identified by MACS2 was performed using HOMER. The top-scoring “known” motif for each time point is shown.

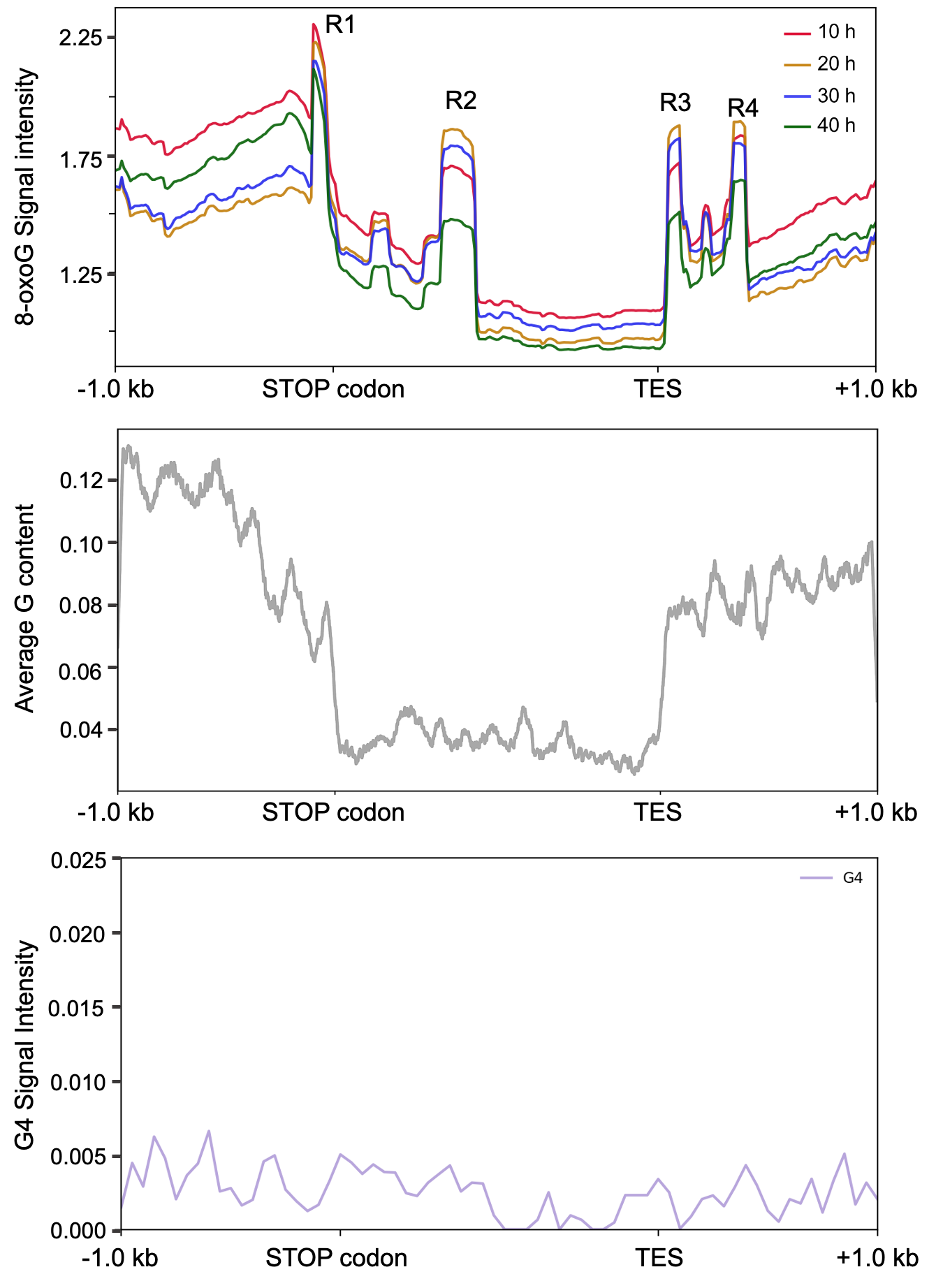

**Supplementary Figure S5:** **Metagene and sequence feature profiles of 3’ends of 8-oxoG-enriched *P. falciparum* genes.** ***Upper panel:*** Metagene profiles showing the average distribution of 8-oxoG signal across all modified genes, from 1 kb upstream of the STOP codon to 1 kb downstream of the Transcription End Site (TES). R1, R2, R3 and R4 indicate highly enriched peaks in this region. ***Middle panel:*** Average G-content (%) across the same genomic regions. ***Lower panel:*** Average density of PQSs obtained from Gazanion et al., 2020 across the same genomics regions.

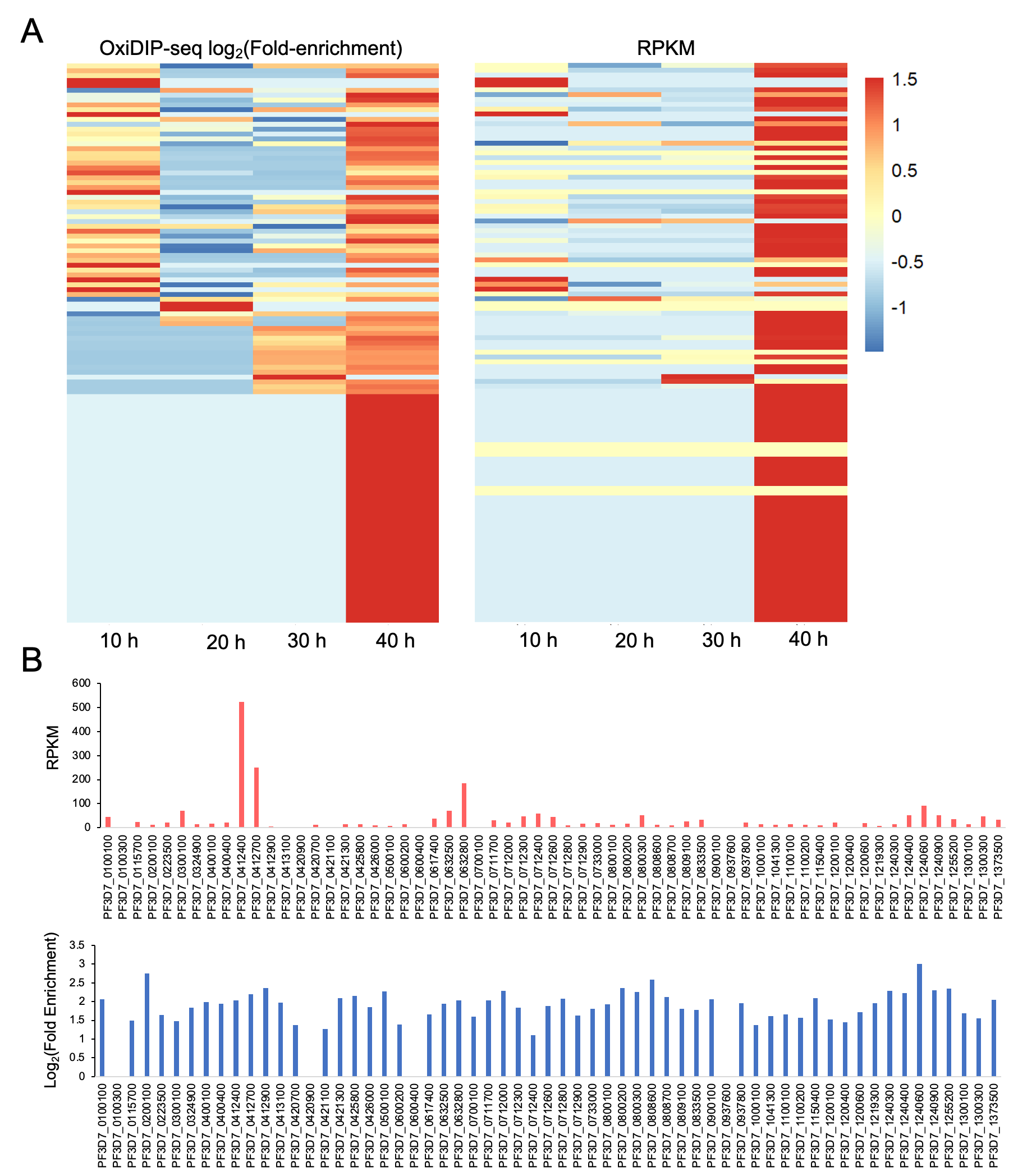

**Supplementary Figure S6: 8-oxoG genic enrichment is not associated with multigene family expression.** **(A)** Heatmaps showing the hierarchical clustering of all rifin genes based on log-transformed and z-normalised 8-oxoG fold enrichment across four IDC stages (***left panel***) and the corresponding log-transformed and z-normalised steady-state transcript levels for the same genes using the 10 h, 20 h, 30 h and 40 h RNA-seq datasets of Toenhake et al., 2018 (***right panel***). Each row represents an individual gene and columns correspond to the indicated time points. The color scale indicates normalized signal intensity (z-score), ranging from low (blue) to high (red). **(B)** ***Upper panel:*** Bar plots showing the expression levels of all *var* genes using RPKM values derived from in-house RNA-seq data generated at 10 hpi (**Supplementary Table S4**). ***Lower panel:*** Bar plots showing the corresponding log_2_FE values of 8-oxoG enrichment from OxiDIP-seq for all *var* genes.

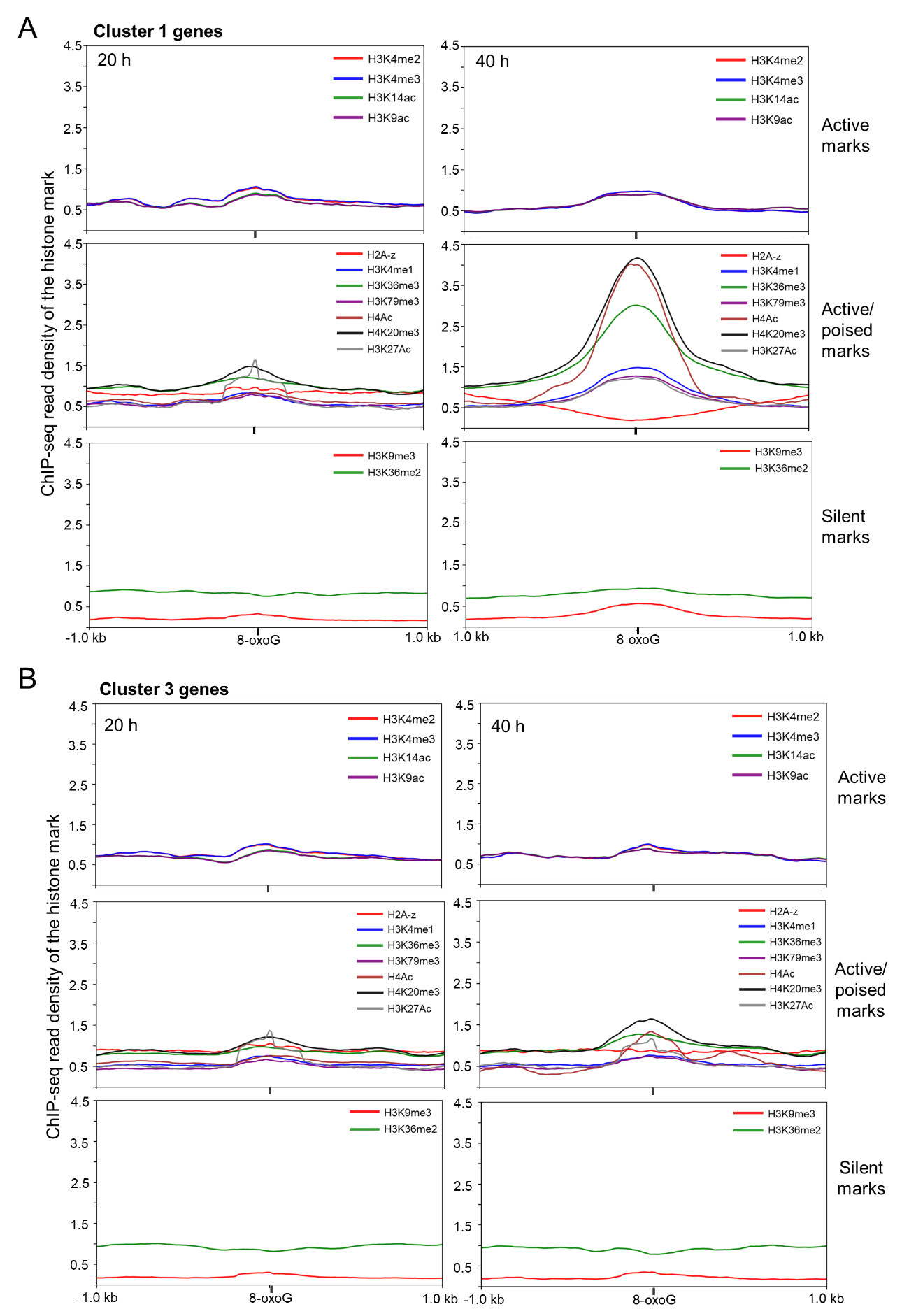

**Supplementary Figure S7:** **Histone post-translational modification (PTM) profiles across 8-oxoG-enriched genes from clusters 1 and 3.** DeepTools plotprofile was used to compare the read density distribution of the indicated active, active/poised and silencing histone PTMs for all 8-oxoG genomic positions located within the exons of modified genes belonging to clusters **(A)** 1 and **(B)** 3 of Figure 4B at 20 h (**left panel**) or 40 h (**right panel**). Profiles represent normalized signal intensity of the histone PTM across the indicated regions. Publicly available ChIP-seq datasets were used for this analysis and include Bartfai et al., 2010, Jiang et al., 2013 and Karmodiya et al., 2015.

**References:**

Bártfai, R., Hoeijmakers, W. A. M., Salcedo-Amaya, A. M., Smits, A. H., Janssen-Megens, E., Kaan, A., Treeck, M., Gilberger, T.-W., Françoijs, K.-J., & Stunnenberg, H. G. (2010). H2A.Z demarcates intergenic regions of the plasmodium falciparum epigenome that are dynamically marked by H3K9ac and H3K4me3. *PLoS Pathogens*, *6*(12), e1001223. https://doi.org/10.1371/journal.ppat.1001223

Gazanion, E., Lacroix, L., Alberti, P., Gurung, P., Wein, S., Cheng, M., Mergny, J.-L., Gomes, A. R., & Lopez-Rubio, J.-J. (2020). Genome wide distribution of G-quadruplexes and their impact on gene expression in malaria parasites. *PLoS Genetics*, *16*(7), e1008917. https://doi.org/10.1371/journal.pgen.1008917

Gorini, F., Scala, G., Ambrosio, S., Majello, B., & Amente, S. (2022). OxiDIP-Seq for Genome-wide Mapping of Damaged DNA Containing 8-Oxo-2’-Deoxyguanosine. *Bio-Protocol*, *12*(21), e4540. https://doi.org/10.21769/BioProtoc.4540

Jiang, L., Mu, J., Zhang, Q., Ni, T., Srinivasan, P., Rayavara, K., Yang, W., Turner, L., Lavstsen, T., Theander, T. G., Peng, W., Wei, G., Jing, Q., Wakabayashi, Y., Bansal, A., Luo, Y., Ribeiro, J. M. C., Scherf, A., Aravind, L., … Miller, L. H. (2013). PfSETvs methylation of histone H3K36 represses virulence genes in Plasmodium falciparum. *Nature*, *499*(7457), 223–227. https://doi.org/10.1038/nature12361

Karmodiya, K., Pradhan, S. J., Joshi, B., Jangid, R., Reddy, P. C., & Galande, S. (2015). A comprehensive epigenome map of Plasmodium falciparum reveals unique mechanisms of transcriptional regulation and identifies H3K36me2 as a global mark of gene suppression. *Epigenetics & Chromatin*, *8*, 32. https://doi.org/10.1186/s13072-015-0029-1

Toenhake, C. G., Fraschka, S. A.-K., Vijayabaskar, M. S., Westhead, D. R., van Heeringen, S. J., & Bártfai, R. (2018). Chromatin Accessibility-Based Characterization of the Gene Regulatory Network Underlying Plasmodium falciparum Blood-Stage Development. *Cell Host & Microbe*, *23*(4), 557-569.e9. https://doi.org/10.1016/j.chom.2018.03.007
